# Inhalable polymer nanoparticles for versatile mRNA delivery and mucosal vaccination

**DOI:** 10.1101/2022.03.22.485401

**Authors:** Alexandra Suberi, Molly K. Grun, Tianyang Mao, Benjamin Israelow, Melanie Reschke, Julian Grundler, Laiba Akhtar, Teresa Lee, Kwangsoo Shin, Alexandra S. Piotrowski-Daspit, Robert J. Homer, Akiko Iwasaki, Hee Won Suh, W. Mark Saltzman

## Abstract

An inhalable platform for mRNA therapeutics would enable minimally invasive and lung targeted delivery for a host of pulmonary diseases. Development of lung targeted mRNA therapeutics has been limited by poor transfection efficiency and risk of vehicle-induced pathology. Here we report an inhalable polymer-based vehicle for delivery of therapeutic mRNAs to the lung. We optimized biodegradable poly(amine-co-ester) polyplexes for mRNA delivery using end group modifications and polyethylene glycol. Our polyplexes achieved high transfection of mRNA throughout the lung, particularly in epithelial and antigen-presenting cells. We applied this technology to develop a mucosal vaccine for SARS-CoV-2. Intranasal vaccination with spike protein mRNA polyplexes induced potent cellular and humoral adaptive immunity and protected K18-hACE2 mice from lethal viral challenge.

**One-sentence summary:** Inhaled polymer nanoparticles (NPs) achieve high mRNA expression in the lung and induce protective immunity against SARS-CoV-2.

## Main Text

mRNA-based vaccines for SARS-CoV-2 have demonstrated the enormous potential of mRNA therapeutics for safe and effective use in the general population (1, 2). The long-anticipated development of mRNA vaccines was enabled by critical advancements in mRNA technology that improved stability and transfection while minimizing innate immune activation (3). To capitalize on these advancements, and expand the application of mRNA therapeutics beyond delivery of systemically administered vaccines, further research and development is required to optimize mRNA delivery vehicles for diverse applications *in vivo* (4, 5). In particular, inhalable mRNA delivery vehicles would enable minimally invasive and lung targeted therapies for pulmonary diseases. Inhaled delivery would be ideal for creating improved mucosal vaccines for respiratory pathogens, protein supplementation, or gene editing in the lung (6-9). However, the lipid nanoparticle (LNP) materials that are currently used for intramuscular mRNA SARS-CoV-2 vaccines are not easily adapted for inhalation delivery (10).

Several major limitations have hindered the development of inhaled mRNA therapeutics. First, mRNA delivery vehicle efficacy is highly dependent on the route of administration (11) and must therefore be optimized for expression in the lung. High transfection efficiency is required to reduce the therapeutic dose and reach the concentration of protein necessary to achieve a therapeutic response. For example, despite initially promising safety and tolerability, RESTORE-CF—an ongoing inhaled mRNA clinical trial, delivering CFTR mRNA in LNPs for treatment of cystic fibrosis— failed to improve pulmonary function in the second interim report, highlighting the need for improvements in the delivery vehicle (12). Another key concern for inhaled therapeutic delivery is that the respiratory mucosa is particularly susceptible to immunopathology (13). Several components of the LNP delivery vehicles used in both approved mRNA vaccines have been shown to induce inflammation in the respiratory tract after intranasal administration (10). A NP optimized for inhaled administration that achieves high mRNA transfection efficiency without inducing an inflammatory immune response that damages the lungs, is needed to enable development of pulmonary mRNA therapeutics.

We sought to overcome the challenges to delivering inhaled mRNA by creating an inhalable, non-inflammatory, polymer-based delivery vehicle. Previously, we have demonstrated that a family of nontoxic biodegradable poly(amine-co-ester) (PACE) polymers can encapsulate and protect nucleic acid cargos for delivery *in vivo* (14). The chemical composition of PACE polymers is highly tunable depending on the monomer components added to the polymerization reaction, the ratios of the components, and synthesis conditions (15). For certain polymer compositions, PACE polymers form polyplexes with mRNA (PACE-mRNA) through a combination of electrostatic interactions between the mildly cationic polymer and the negatively charged phosphate backbone of nucleic acids as well as hydrophobic interactions between segments of the polymer chain. PACE polymers can also be modified through the addition of end groups. We have demonstrated that amine containing end groups can improve transfection efficiency by facilitating endosomal escape of mRNA from the endocytosed NP into the cytoplasm (16). Stabilization of PACE-mRNA polyplexes with polyethylene glycol (PEG) can further improve mRNA delivery *in vivo* (17). We capitalized on the highly tunable nature of PACE polyplexes by screening a library of delivery vehicles with different chemical end groups and PEG contents to optimize for high protein expression after local delivery to the respiratory tract.

In the present work, we created an optimized PACE-mRNA polyplex which achieves high protein expression in the lung— primarily in epithelial cells and antigen presenting cells— without inducing deleterious inflammatory responses. To demonstrate the translational potential of our delivery vehicle, we created an inhalable spike (S) protein mRNA vaccine for SARS-CoV-2. Our vaccine induced *de novo* immunity to SARS-CoV-2 through both systemic and local induction of antibodies. We demonstrated effective draining lymph node germinal center activation, resulting in expansion of S specific memory B cells and antibody secreting cells. Inhaled vaccination induced circulating antigen specific CD8^+^ T cells and lung resident S specific tissue memory CD8^+^ T cells. Finally, intranasal PACE-mRNA vaccination protected K18-hACE2 mice from lethal viral challenge. This work represents the first report of an inhaled nonviral, mRNA-based vaccine capable of eliciting *de novo* immunity against SARS-CoV-2 without the need for intramuscular priming.

## Results

### Characterization of PACE-mRNA polyplexes

We optimized PACE polyplexes for mRNA delivery to the lung by screening novel blends that combine the benefits of PEG stabilization and end group mediated enhanced endosomal escape. We identified 10 promising end group chemistries and synthesized PACE polymers by enzymatic copolymerization of 15-pentadecanolide (PDL), N-methyl diethanolamine (MDEA), and sebacic acid (SA) followed by carbodiimidazole (CDI)-mediated conjugation of amine containing end groups (Fig. 1A, S1-4) (16). A PACE block co-polymer (PACE-PEG) was synthesized by adding a 5 kDa methoxy-ended PEG (mPEG) to the reaction mixture, in a modified literature protocol (15, 18) (table S1).

**Fig 1.**
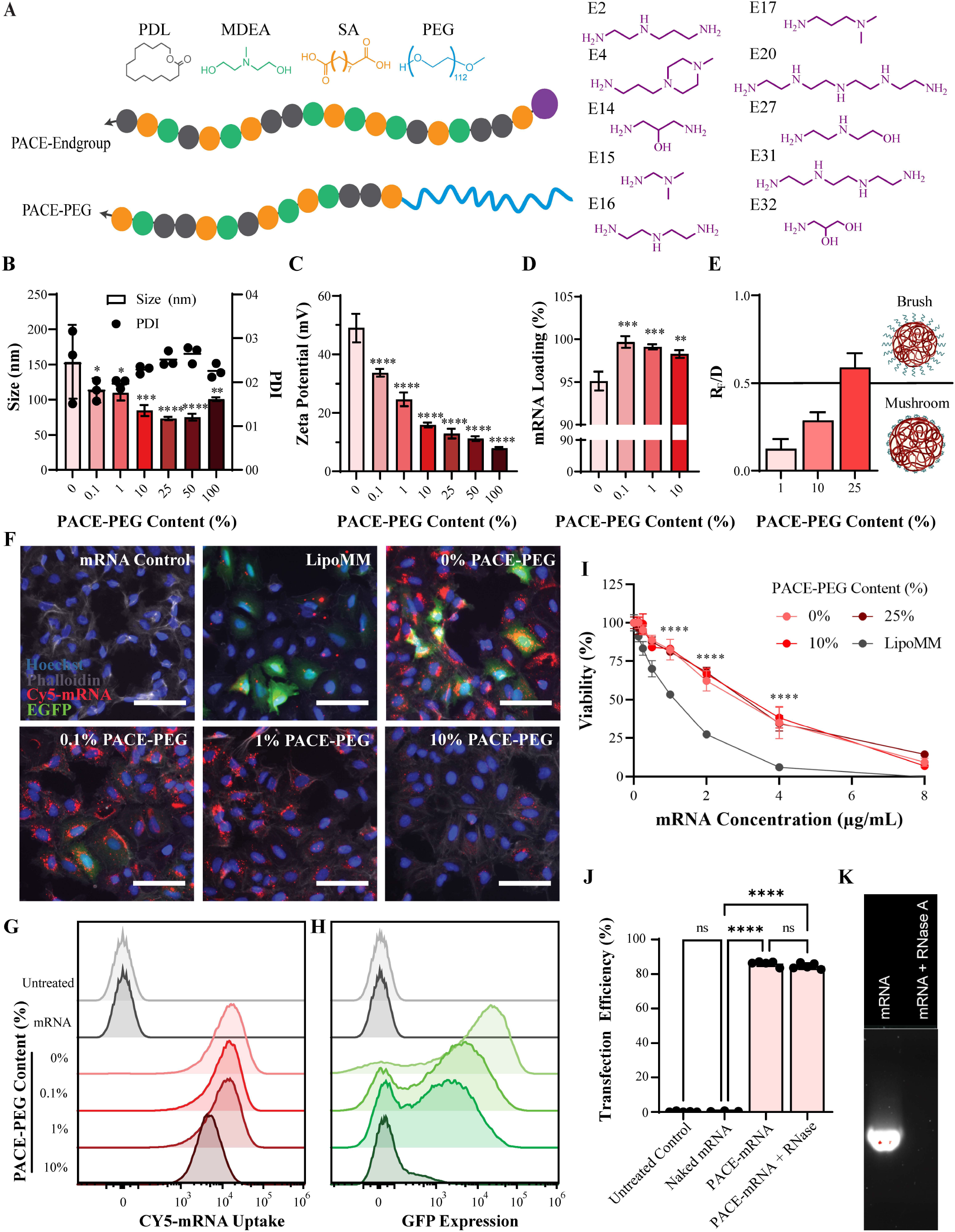
Characterization of PACE-mRNA polyplexes and *in vitro* activity. (**A**) Schematic of end group-modified and PEGylated PACE polymer composition with chemical structures of base monomers and end groups (**B**) Size and PDI; (**C**) zeta potential; (**D**) mRNA loading; and (**E**) PEG conformation on the surface of PACE-E14 polyplexes with varying PACE-PEG content. *R*_f_ represents PEG Flory radius and *D* the distance between PEG chains. Asterisks indicate statistical difference from non-PEGylated polyplexes. (**F**) Representative mRNA uptake and transfection efficiency of Cy5-conjugated EGFP mRNA delivered with PACE-PEG blended polyplexes (scale bar, 75 μm). (**G**) Uptake of Cy5-conjugated mRNA and (**H**) transfection efficiency of EGFP mRNA in A549 cells with PEGylated PACE-E14. (**I**) *In vitro* cytotoxicity of PEGylated PACE-E14 polyplexes compared to Lipofectamine MessengerMAX. Asterisks indicate differences between all PACE polyplexes and Lipofectamine. (**J-K**) Transfection efficiency of EGFP mRNA PACE-E14 polyplexes in HEK293T cells with or without coincubation of RNase and a gel run with either naked mRNA or mRNA and RNase showing degradation of mRNA by the enzyme. * *p* ≤ 0.05, ** *p* ≤ 0.01, *** *p* ≤ 0.001, **** *p* ≤ 0.0001. Data are pooled from two independent experiments.

We administered end group modified PACE-mRNA to human alveolar epithelial cells (A549) and identified PACE with end group 14 (E14) as a promising candidate for delivery (fig. S5). Next, we examined the effect of PACE-PEG content on polyplex characteristics by blending PACE-E14 and PACE-PEG at different ratios. PACE-PEG reduced the size and surface charge of PACE-mRNA polyplexes (Fig. 1B-C), which is consistent with previous reports that PEGylation of cationic polymers can neutralize surface potential (19-21). mRNA loading efficiency was measured using a fluorescent nucleic acid stain (Quant-iT RiboGreen). Although non-PEGylated polyplexes efficiently loaded mRNA (95% loading), the addition of PACE-PEG improved loading efficiency to around 99% (Fig. 1D), suggesting that incorporating PEG can improve the polyplex formulation process.

To provide greater insight into how PEG content affects polyplex structure, we calculated the PEG density and conformation on the surface of PACE-mRNA polyplexes with 1%, 10%, or 25% PACE-PEG by first determining the polyplex molar mass through static light scattering (SLS) measurements (22). We calculated the PEG density on the surface of polyplexes (table S2). Polyplexes with PACE-PEG content of 1-10% (by weight) possessed a surface PEG density in which PEG chains are in a mushroom conformation, whereas 25% PACE-PEG content produces a surface PEG density corresponding to a brush conformation (Fig. 1E).

To compare the mRNA uptake and transfection efficiency of PACE-mRNA polyplexes *in vitro*, we treated A549 cells with polyplexes loaded with Cy5-labeled EGFP mRNA or unlabeled EGFP mRNA using a range of PACE-PEG contents, and we measured the fluorescent signal by microscopy and flow cytometry. Fluorescence microscopy demonstrated that PACE-mRNA without PACE-PEG transfected cells at a comparable rate to a commercial transfection reagent, Lipofectamine MessengerMax (LipoMM). However, we observed a noticeable decrease in EGFP expression at 0.1% and 1% PACE-PEG while the Cy5-labeled mRNA signal remained consistent (Fig. 1F). By flow cytometry, we observed that although polyplex uptake (measured by Cy5-mRNA signal) decreased with the addition of PACE-PEG, the overall uptake was still high; even at 10% PACE-PEG concentration, 99.7% of cells were positive for Cy5 (Fig. 1G). By contrast, the average transfection efficiency (measured by EGFP signal) decreased from 85% to 15% with increasing amounts of PACE-PEG (Fig. 1H). Overall, the incorporation of PACE-PEG caused a decrease in mRNA transfection efficiency that outpaced the drop in mRNA uptake (fig. S6). These results suggest that—while PEGylation inhibits the uptake of PACE polyplexes to some degree—lower uptake doesn’t fully explain the inhibitory effect of PEG on mRNA transfection *in vitro*. Based on our prior work demonstrating that mRNA transfection efficiency correlates more strongly with endosomal escape than cellular uptake (16), we hypothesize that PEGylation influences the extent of endosomal escape. In support of this, we noted a decrease in cytoplasmic Cy5-mRNA signal as the extent of PEGylation was increased (Fig. 1F).

The biocompatibility of PACE-mRNA polyplexes with 0%, 10%, and 25% PACE-PEG was compared to LipoMM using the neutral red viability assay. A549 cells were treated with a range of mRNA concentrations, delivered with a consistent ratio of polymer or LipoMM. At higher concentrations, the PACE-mRNA polyplexes were significantly less toxic than LipoMM. There was no significant difference in viability between the polyplexes based on PEG content (Fig. 1I). These results demonstrate the superiority of PACE-mRNA polyplexes compared to LipoMM in achieving comparably high mRNA transfection efficiency with reduced cytotoxicity *in vitro*.

A primary motivation for encapsulating mRNA into a delivery vehicle is to protect the delicate nucleic acid from abundant degradative enzymes in the lung mucosa. To demonstrate the ability of PACE to protect cargo from RNase activity, we co-incubated PACE-EGFP mRNA polyplexes with RNase before treating HEK293 cells. RNase pretreatment had no significant effect on transfection efficiency as measured by flow cytometry analysis of the EGFP signal (Fig. 1J). PACE preserved the mRNA integrity for delivery to cells, whereas naked mRNA was completely degraded after incubation with RNase under identical conditions (Fig. 1K).

### *In vivo* optimization of PACE-mRNA polyplexes

Prior research has established that traditional cell culture methods are not predictive of transfection *in vivo* for both viral and non-viral delivery systems (5, 17, 23-25). PEGylation of NPs presents additional advantages with administration to mucosal surfaces, as PEGylation is well known to facilitate transport through mucus (26), which is not present in most cell cultures. Therefore, despite the reduced transfection efficiency of PEGylated polyplexes observed *in vitro*, we performed an *in vivo* screen with PACE-PEG content ranging from 0% (no PACE-PEG) to 100% (no PACE-E14). We delivered 5 µg of firefly luciferase (FLuc) mRNA to mice by intratracheal instillation (IT) to assess pulmonary transfection efficiency via a readout of luminescence. At 24 hours, non-PEGylated polyplexes and polyplexes containing up to 50% PACE-PEG had a strong luminescent signal that was 1,000x – 10,000x higher than untreated animals (Fig. 2A). Low amounts of PACE-PEG incorporation (1%, 10%, 25%) improved transfection efficiency compared to non-PEGylated polyplexes, and we observed the highest luminescence with 10% PACE-PEG. With higher levels of PACE-PEG (50%, 100%), there was a significant decrease in luciferase expression, and polyplexes made entirely with PACE-PEG (100%) yielded no luciferase signal. All PACE-mRNA polyplexes containing PACE-E14 content outperformed a commercially available transfection agent, jetRNA. We confirmed these results by *In Vivo* Imaging System (IVIS) imaging, where we observed the highest luminescent signal using 10% PACE-PEG polyplexes (Fig. 2B). We also tracked the luminescent signal after a single delivery over time to see if PEGylation altered the kinetics of mRNA delivery (fig. S7). Luciferase expression was highest at 6 and 24 hours, which is consistent with previous reports of mRNA expression in the lung (11, 27). At 6 and 24 hours, significantly higher luminescence was observed in the group treated with 10% PACE-PEG polyplexes compared to non-PEGylated polyplexes, but at later time points, there was no significant difference between experimental groups. These results demonstrate that low levels of vehicle PEGylation can improve delivery and translation of mRNA, which is consistent with our previous report of mRNA delivery with PACE polyplexes that were not end group-modified (17); however, here we report significantly higher transfection levels—over 100-fold higher by luminescence—demonstrating the significant benefit of using end group-modified (E14) PACE.

**Fig 2.**
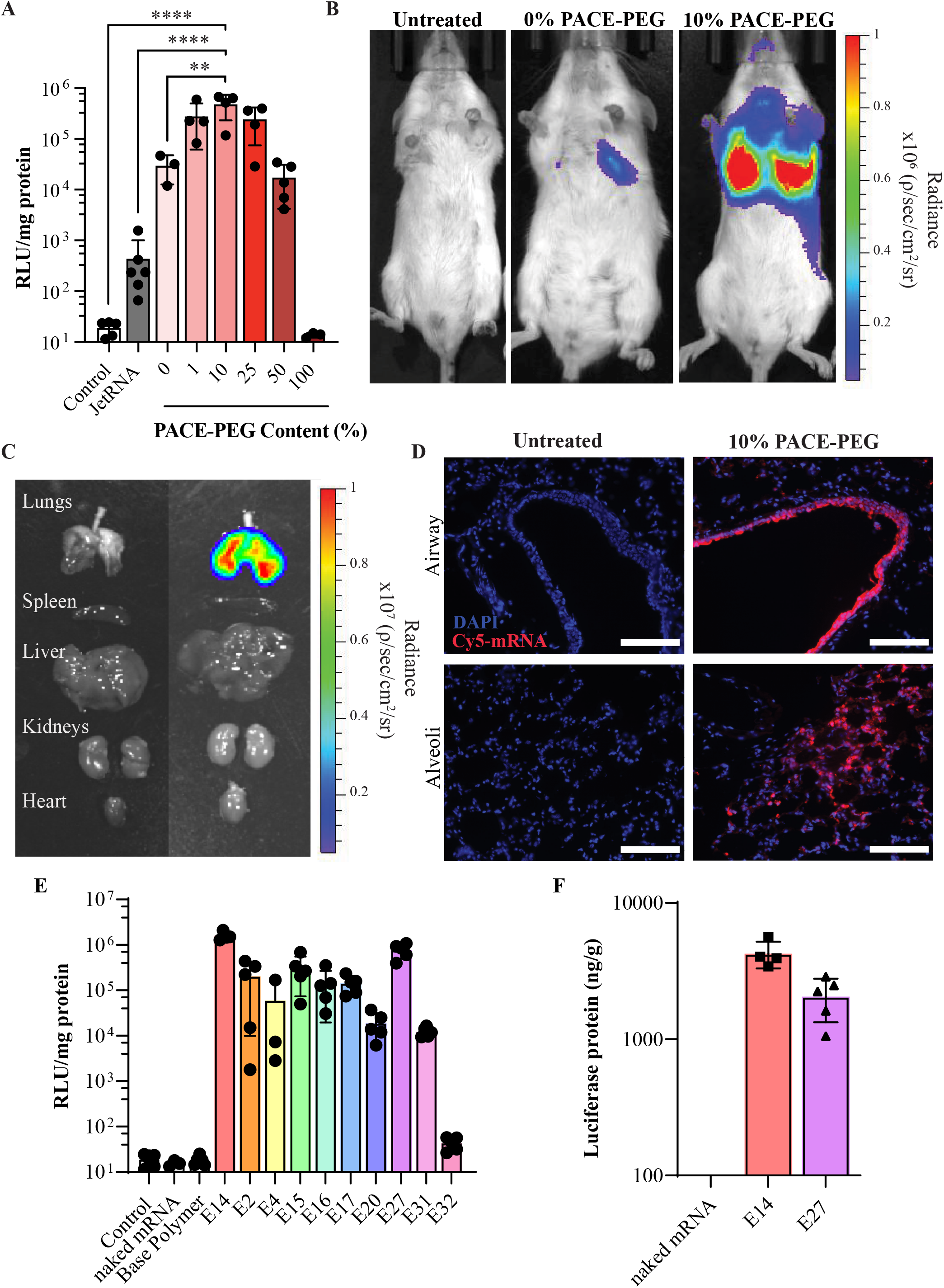
IT mRNA delivery with PACE-mRNA polyplexes provides significant protein expression and mRNA distribution throughout the airways and parenchyma. (**A**) Luciferase protein expression in lung tissue after delivery of FLuc mRNA with PEGylated PACE-E14 and *in vivo*-jetRNA. Sample means of log-transformed values were compared by Tukey’s multiple comparison test. (**B**) Representative luciferase expression by IVIS 24 hours after IT delivery of PACE-E14 10% PACE-PEG FLuc mRNA polyplexes in animals and (**C**) in explanted organs. (**D**) Distribution of Cy5-conjugated mRNA in the lung 30 minutes after delivery with PACE-E14 polyplexes (scale bar, 150 μm). (**E**) Luciferase protein expression in lung tissue after delivery of FLuc mRNA with polyplexes of various end group modified and 10% PACE-PEG content, untreated control, naked mRNA control, and base polymer with no end group control. (**F**) Quantification of luciferase protein extracted from lungs 24 hours after treatment with either E14 or E27 polyplexes with 10% PACE-PEG. * *p* ≤ 0.05, ** *p* ≤ 0.01, *** *p* ≤ 0.001, **** *p* ≤ 0.0001. Data are pooled from two independent experiments.

Next, we performed IVIS imaging of explanted organs (lungs, spleen, liver, kidneys, and heart), which confirmed that luciferase expression was entirely localized in the lungs, with no observable luminescence coming from other organs (Fig. 2C). High luminescent signal was achieved in both left and right sided lobes of the lung. Using Cy5-labeled mRNA to assess the distribution of IT delivered PACE-mRNA polyplexes (E14 with 10% PACE-PEG) within the pulmonary architecture, we found PACE-mRNA throughout the large airways and into the alveolar regions of the lung parenchyma (Fig. 2D).

While we had found that E14 modified PACE led to optimal *in vitro* epithelial cell transfection efficiency, we wondered if this would hold *in vivo*. To test the *in vivo* efficacy of NPs formed from different end group-modified versions of PACE, we performed an additional screen with all 10 end group-modified PACE polymers (Fig. 1A) and a fixed PACE-PEG content of 10% by weight (Fig. 2E). No measurable luminescent signal was observed following administration of 5 µg of naked mRNA in buffer or with polyplexes produced using the base polymer that did not have a conjugated end group. All end group-modified PACE-mRNA polyplexes achieved significant mRNA expression above control except for E32. We found that E14 remained the top performing end group for transfection in the lung, consistent with our preliminary cell culture-based screen. However, several other end groups also achieved high protein expression. E27 was identified as a second promising end group for PACE-mRNA polyplexes. We further quantified the amount of luciferase protein per gram of total protein in samples extracted from homogenized lungs by comparing to a standard curve of luminescence with recombinant firefly luciferase protein. Quantification of the two formulations with highest mRNA transfection (E14 and E27 with 10% PACE-PEG) each contained more than 1000 ng of FLuc per gram protein in the lung (Fig. 2F).

### PACE-mRNA polyplexes transfect epithelial cells and antigen presenting cells in the lung

Having identified two end group-modified PACE polymers (E14 and E27) and the optimal PACE-PEG concentration (10%) for achieving high transfection targeted to the lung, we next characterized cell type-specific mRNA expression in the lung using Ai14 tdTomato reporter mice. Ai14 mice have a loxP-flanked STOP cassette upstream of a tdTomato gene, which can be excised by Cre-mediated recombination, enabling tdTomato expression in the cell (Fig. 3A). After IT instillation of 10 µg of Cre mRNA in PACE-mRNA polyplexes (E14 with 10% PACE-PEG), 9.97% (SD 2.32) of all cells in the lung and 28.8% (SD 12.8) of cells in the bronchoalveolar lavage fluid (BALF) expressed tdTomato (Fig. 3B). We further evaluated the lungs to identify endothelial (CD31^+^), epithelial (EpCAM^+^), and leukocyte (CD45^+^) subpopulations (fig. S8). Transfection was predominantly achieved in epithelial cells and leukocytes, with 21.5% (SD 7.32) of lung epithelial cells and 19.6% (SD 3.47) of lung leukocytes expressing tdTomato. Endothelial cells were not significantly transfected (Fig. 3C). Of the leukocyte cells, we further evaluated markers for antigen presenting cells (APCs) in lung tissue. We found that 59.6% (SD 9.43) of CD11c^+^CD11b^+^ cells and 1.04% (SD 0.28) of CD11c^+^CD11b^-^ cells expressed tdTomato. Fluorescence microscopy of lung sections demonstrated that expression of tdTomato could be found primarily in cells lining the conducting airway and throughout the alveolar regions (Fig. 3E). The localization of fluorescence within the lung architecture is consistent with our flow cytometry finding that transfection occurs primarily in epithelial cells and APCs. We found a similar pattern of expression following delivery of E27 polyplexes. Epithelial cells (16.6% SD 3.83) and antigen presenting CD11c^+^CD11b^+^ cells (58.2% SD 7.34) were most highly transfected. However, total lung cell transfection was slightly reduced (7.37% SD 2.69) (fig. S9).

**Fig 3.**
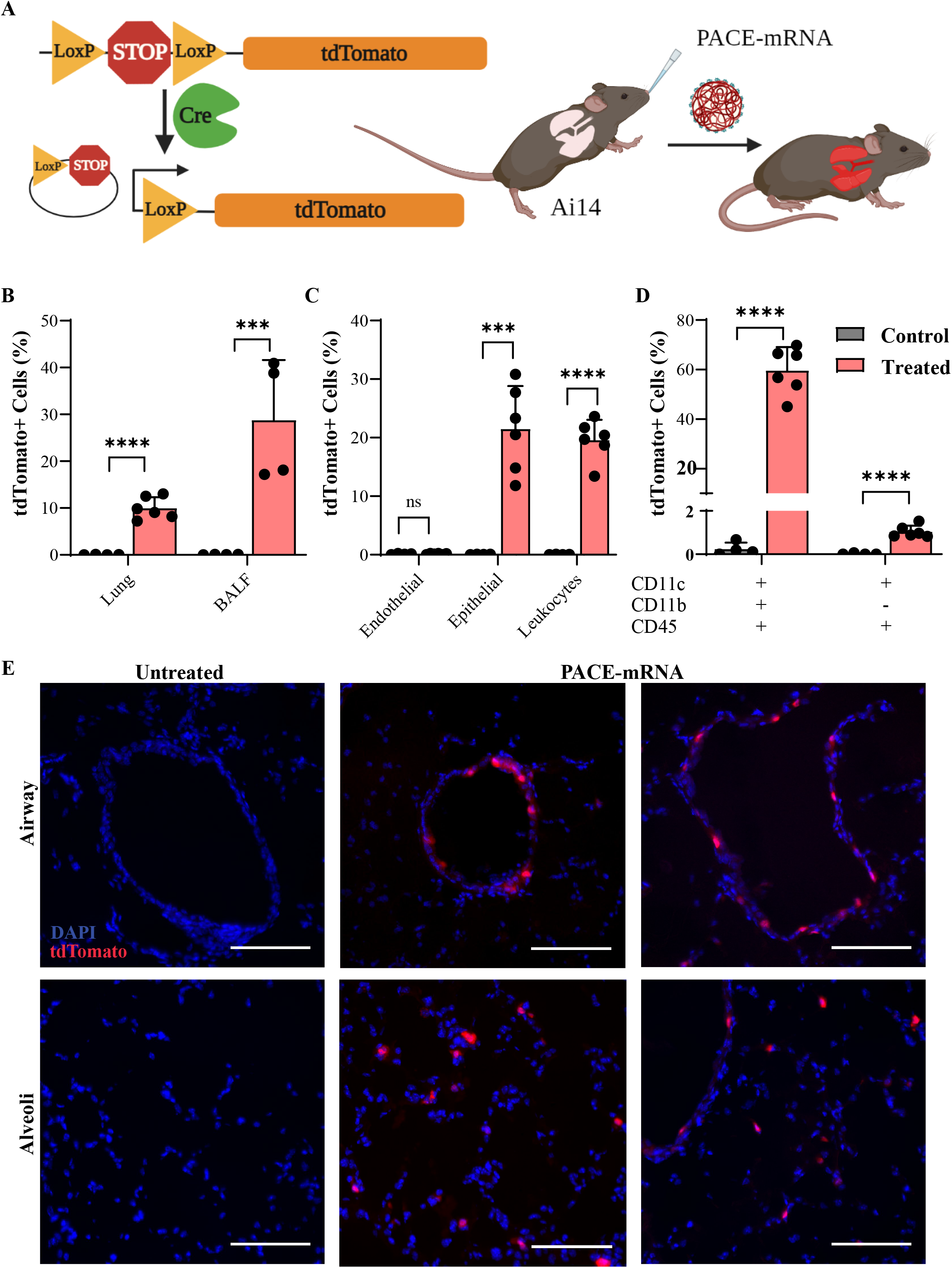
mRNA expression after IT PACE-mRNA delivery occurs in epithelial cells and APCs. (**A**) Schematic of Cre mediated recombination in Ai14 mice resulting in expression of tdTomato protein in transfected cells. Percent of (**B**) all live cells in the lung and BALF, (**C**) endothelial, epithelial, or leukocyte cells in the lung and (**D**) APCs in the lung that express tdTomato 24 hours after administration of PACE-E14 polyplexes (10% PACE-PEG) loaded with Cre mRNA. Statistical significance was calculated by multiple unpaired t-test with holm-sidak method. (**E**) Representative images of control (untreated) and PACE-E14 polyplex (10% PACE-PEG) treated lungs from Ai14 mice by fluorescence microscopy (scale bar, 100 μm, DAPI in blue, tdTomato in red). * *p* ≤ 0.05, ** *p* ≤ 0.01, *** *p* ≤ 0.001, **** *p* ≤ 0.0001. Data are pooled from two independent experiments.

The *in vivo* biocompatibility of PACE-E14 formulations (with 10% PACE-PEG) was compared to buffer-only treated lungs after 48 hours. Analysis was performed by a pathologist blinded to the treatment group. Histology of treated lungs showed some focal areas with mild neutrophilic infiltrate in the terminal airways only. No evidence of necrosis or other acute airway epithelial change was present (fig. S10).

### Intranasal PACE-mRNA vaccination effectively induces antigen specific B and T cell adaptive immune responses in the mediastinal lymph node

Next, we sought to assess whether our top performing PACE-mRNA polyplex (E14 with 10% PACE-PEG) could be applied therapeutically as an inhaled vaccine. We encapsulated mRNA encoding the spike (S) protein from SARS-CoV-2 into PACE-mRNA polyplexes. We chose a mouse model, K18-hACE2 mice which express human ACE2 (the entry receptor for SARS-CoV-2) from the mouse cytokeratin 18 promoter. K18-hACE2 mice are commonly used in preclinical studies due to their susceptibility to infection and severe pulmonary disease following SARS-CoV-2 viral challenge (28). We used a prime and boost vaccination strategy in which mice received a 10 µg dose of PACE-mRNA delivered intranasally on days 0 and 28. On day 42 (14 days post-boost), we assessed for the development of adaptive immunity against the S protein (Fig. 4A).

**Fig 4.**
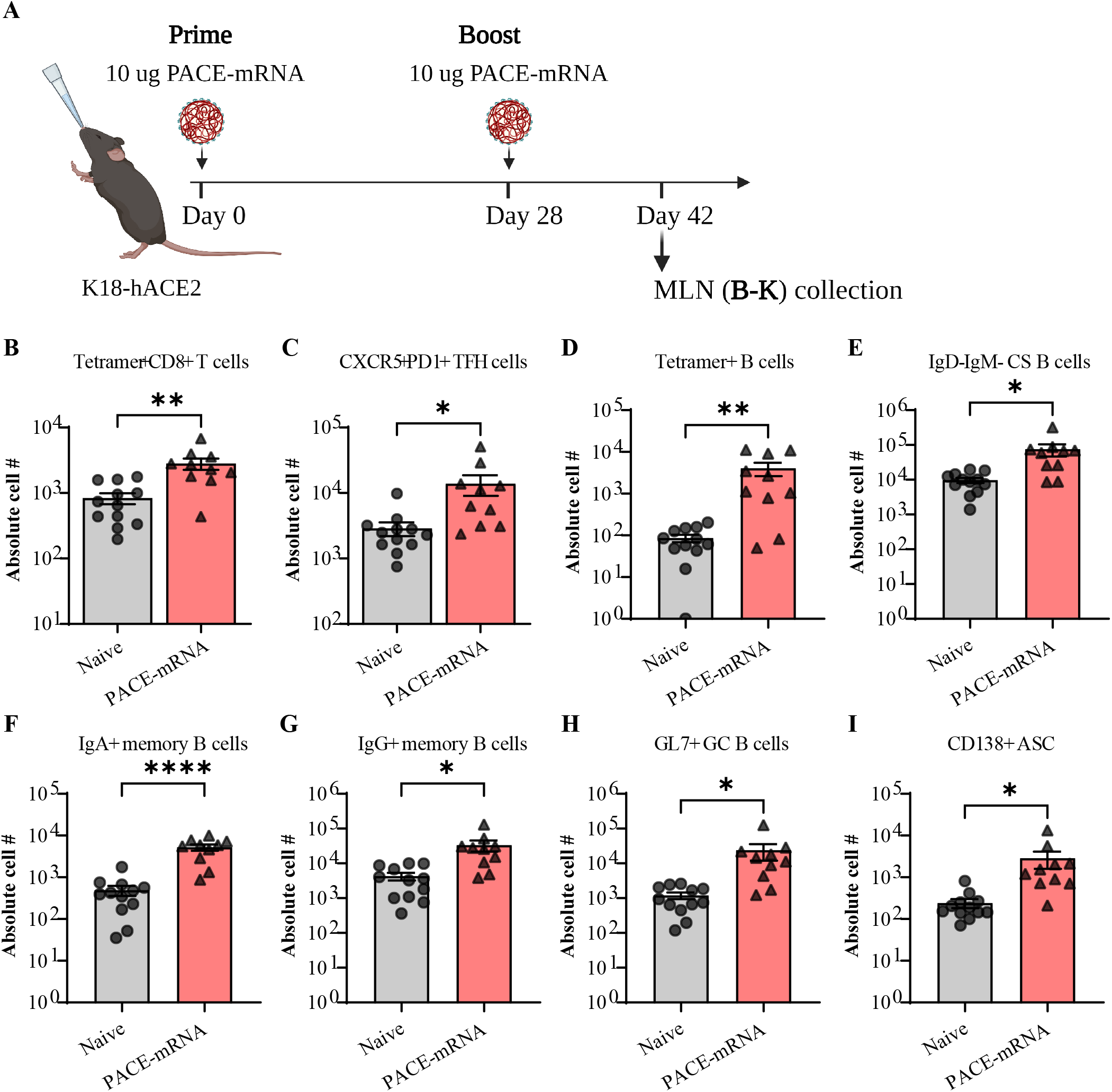
IN PACE-mRNA vaccination induces antigen specific T and B cell responses in the draining lymph node. (**A**) Schematic of PACE-mRNA vaccination in K18-hACE2 mice. Mice were primed (day 0) and boosted (day 28) with a 10 µg dose of S protein mRNA encapsulated in PACE-E14 polyplexes with 10% PACE-PEG. MLN were harvested on day 42 for analysis. (**B**) Quantification of extravascular (IV^-^) SCV2 spike-specific Tetramer^+^ CD8 T cells in MLN. (**C**) Quantification of extravascular (IV^-^) CXCR5^+^PD1^+^ T_FH_ cells in MLN. (**D-I**) Quantification of various extravascular B cell subsets, including RBD tetramer-binding B cells, class switched B cells, IgA^+^ memory B cells, IgG^+^ memory B cells, activated germinal center B cells, and antibody secreting cells in MLN. Mean ± s.e.m.; Statistical significance was calculated by student’s t-test. * *p* ≤ 0.05, ** *p* ≤ 0.01, *** *p* ≤ 0.001, **** *p* ≤ 0.0001. Data are pooled from two independent experiments.

To evaluate for effective PACE-mRNA induction of adaptive immune responses, we assessed the mediastinal lymph node (MLN) by flow cytometry for the development of germinal center responses and antigen specific memory and effector B cells and CD8^+^ T cells, the critical mediators of durable adaptive immunity in the lung (29-31). For identification of S specific T cells, we stained with an MHC class 1 tetramer to a SARS-CoV-2 S protein epitope (VNFNFNGL) and stained B cells with a receptor binding domain (RBD) B cell tetramer. PACE-mRNA vaccination induced a significant population of S specific CD8^+^ T cells (Fig. 4B). Another T cell subtype important for mounting an adaptive immune response is T follicular helper (T_FH_) cells. T_FH_ cells promote affinity maturation and class switch recombination in B cells and critically orchestrate the development of neutralizing antibody responses (32). Systemically administered mRNA vaccines elicit T_FH_ and germinal center B cells, which strongly correlates with neutralizing antibody production (33, 34). We found significant induction of T follicular helper (T_FH_) cells in the MLN (Fig. 4C). Correspondingly, there was significant expansion of S specific B cells (CD19^+^B220^+^Tetramer^+^), class switched B cells expressing IgA and IgG (CD19^+^B220^+^IgD^-^IgM^-^), activated germinal center B cells (CD19^+^B220^+^GL7^+^), and antibody secreting cells (CD138^+^) in the MLN (Fig. 4D-I). These results demonstrated that PACE-mRNA vaccination induces potent antigen specific T and B cell responses in the draining lymph node following mucosal delivery. Further, S specific B cells expressed memory markers demonstrating promise for the efficacy and durability of the adaptive immune response after PACE-mRNA vaccination.

### Intranasal PACE-mRNA vaccination confers protective tissue resident and circulating immunity against SARS-CoV-2 viral challenge

Finally, we assessed lung tissue, serum, and BALF for local and systemic antigen specific T cells and antibodies. We used intravenous (IV) labeling of CD45 to differentiate between circulating and tissue resident immune cells. Additionally, 4 weeks after boost delivery (day 56), we challenged naïve and PACE-mRNA vaccinated K18-hACE2 mice with a lethal dose of SARS-CoV-2 (Fig. 5A). Consistent with our findings in the MLN, we found that vaccination induced a significant population of S specific CD8^+^ T cells in the lung parenchyma (IV^-^) (Fig. 5B). S specific CD8^+^ T cells significantly expressed tissue resident memory (T_RM_) surface markers, CD69^+^ and CD103^+^ (Fig. 5C-D). We also found significant increases in systemic circulating antigen specific CD8^+^ T cells harvested from the lungs (IV^+^Tetramer^+^CD8^+^) (Fig. 5E). These results demonstrate that vaccination elicited both a lung resident and a circulating CD8^+^ T cell response. Currently available intramuscularly administered vaccines induce circulating but not lung resident S specific T_RM_ responses (29), highlighting the advantage of our mucosal PACE-mRNA delivery strategy.

**Fig 5.**
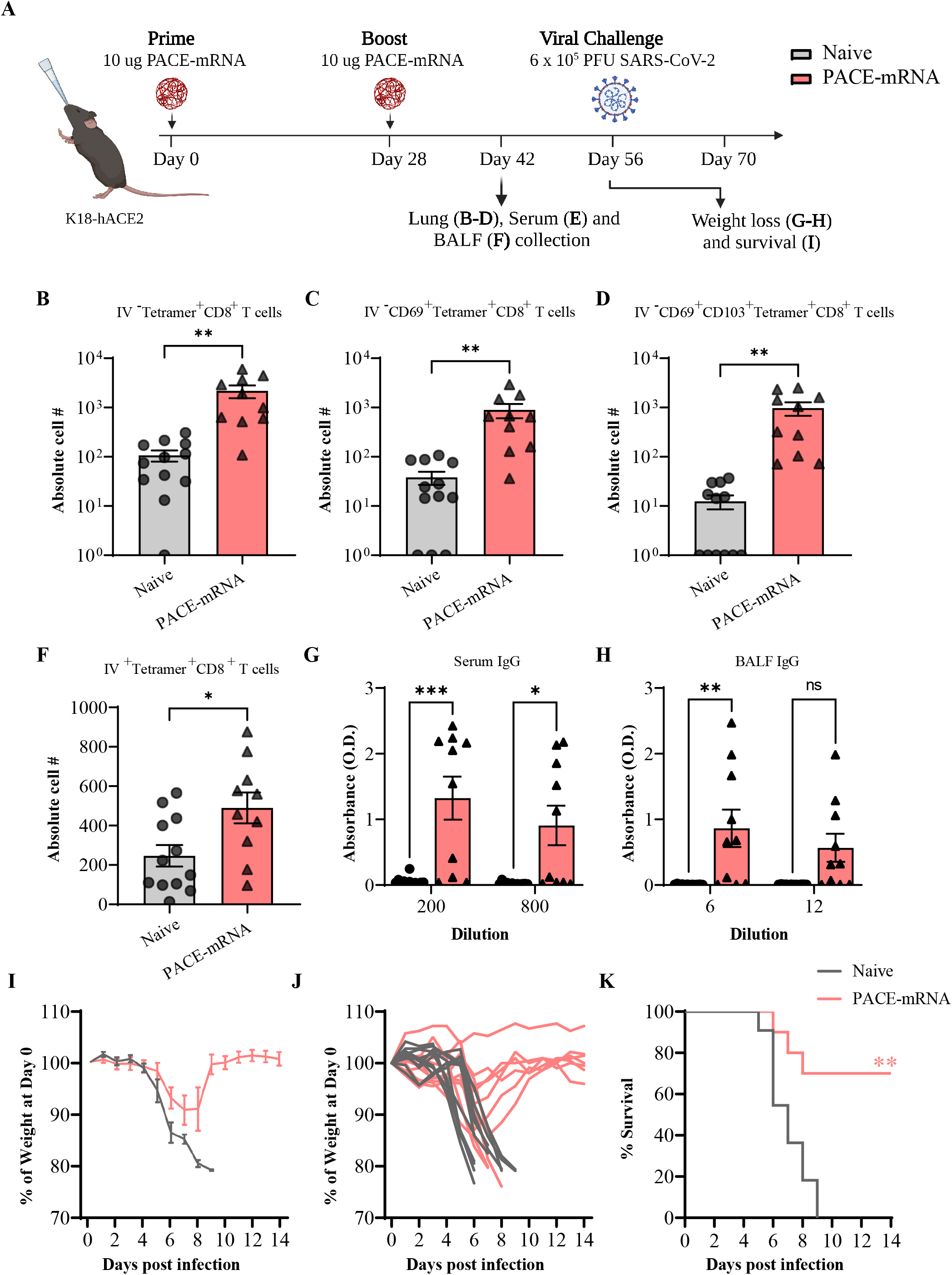
IN PACE-mRNA vaccination induces protective cellular and humoral immunity. (**A**) Schematic of PACE-mRNA vaccination in K18-hACE2 mice. Mice were primed (day 0) and boosted (day 28) with a 10 µg dose of S protein mRNA encapsulated in PACE-E14 polyplexes with 10% PACE-PEG. Lungs, serum, and BALF were harvested on day 42 for analysis. An additional group of vaccinated animals was challenged with 6×10^5^ PFU of SARS-CoV-2 on day 56 and weight loss and survival were evaluated over two weeks compared to untreated naïve mice. (**B-D**) Quantification of extravascular (IV^-^) SCV2 spike-specific Tetramer^+^ CD8 T cells, CD69^+^CD103^-^Tetramer^+^ CD8 T cells, or CD69^+^CD103^+^Tetramer^+^ CD8 T cells in the lung. (**E**) Quantification of circulating (IV^+^) SCV2 spike-specific Tetramer^+^ CD8 T cells from lung vasculature. (**F**) Serum and (**G**) BALF measurement of SCV2 spike S1 subunit-specific IgG. Mean ± s.e.m.; Statistical significance was calculated by student’s t-test. (**H-I**) Average and (**I**) individual weight measurements after viral challenge in nave and PACE-mRNA vaccinated mice. (**J**) Survival of naïve and vaccinated mice from 1 to 14 days post infection. Mean ± s.e.m.; Statistical significance was calculated by log-rank Mantel–Cox test. * *p* ≤ 0.05, ** *p* ≤ 0.01, *** *p* ≤ 0.001, **** *p* ≤ 0.0001. Data are pooled from two independent experiments.

We assessed serum and BALF for anti-SARS-CoV-2 spike S1 IgG and IgA to determine whether mucosal vaccination induced humoral immunity. We found that both circulating (serum) and mucosal (BALF) IgG antibodies were significantly induced (Fig. 5F-G), but IgA was not significantly induced (fig. S11). Finally, to assess whether the demonstrated adaptive immunity would protect mice from viral infection, we challenged mice with a lethal dose (6×10^5^ PFU) of SARS-CoV-2. Weight loss and survival in vaccinated mice were both significantly improved (Fig. 5H-J), demonstrating the protective efficacy of our PACE-mRNA polyplexes.

## Discussion

We have demonstrated that PACE polymer formulations can be optimized and applied for local mRNA delivery to the lung. We employed a screening method for optimizing polyplexes, a strategy which has previously been employed for lipopolyplex (25) and LNP (35-37) optimization. LNP optimization often requires screening varying ratios of four component variables, generating large and unwieldy chemical spaces. Our polyplexes required only two components which facilitated screening efficiency. We showed that a relatively simple polyplex design containing 90% end group-modified PACE (PACE-E14) and 10% PACE-PEG complexed with mRNA to form small and stable polyplexes which protect mRNA from enzymatic degradation and achieve high *in vivo* transfection efficiency.

By calculating the PEG surface density, we gained insight into how PEG content affects polyplex structure. The change in conformation from the more compact mushroom to the brush conformation at 25% PACE-PEG corresponded with the drop in transfection efficiency observed *in vivo* with higher PACE-PEG content. These results are consistent with previous studies demonstrating that PEG surface coverage at high densities can interfere with cellular uptake *in vivo* (17, 23). The transition from the mushroom to the brush conformation results in a thicker hydrophilic barrier, thereby decreasing the potential for interaction between the cell membrane and the PACE end group. This data demonstrates that conformational changes that decrease the exposure of cationic surface elements can interfere with transfection.

This work represents the first polyplexes of end group-modified PACE with PACE-PEG polymers and the first investigation of end group-modified PACE-mRNA in the lung. Previous work in both polymer (38, 39) and lipid (40, 41) design has demonstrated that amine containing end groups can increase mRNA expression or particle targeting to the lung. Our screen identified several amine-containing end groups that increase expression in the lung. The top performing end groups in our screen (E14 and E27) were not the same end groups which transfected most efficiently following intravenous administration of unPEGylated PACE-mRNA polyplexes (16), supporting the need for compartment-specific optimization of delivery vehicles.

Our optimized PACE polyplexes facilitated significant mRNA transfection into epithelial cells and antigen presenting cells. Ease of epithelial cell targeting is a primary advantage of inhaled delivery strategies (27). Protein replacement therapy via mRNA delivery to epithelial cells is therapeutically relevant for a host of diseases such as cystic fibrosis (CF) (42), asthma (43), surfactant B protein deficiency (44), and alpha-1-antitrypsin deficiency (45). Our epithelial transfection rate of >20% after a single dose suggests that protein expression will be therapeutically relevant; prior research has shown that for significant disease mitigation only a fraction of lung cells in CF need to express CFTR. For example, 17-28% of cells expressing CFTR in a porcine lung model restored 50% of normal CFTR function, an amount consistent with significant amelioration of symptoms (46). Previous attempts to deliver inhaled mRNA in preclinical models (47, 48) have achieved moderate success in disease treatment, demonstrating both proof of concept for the therapeutic avenue and the need for further innovation. The significant epithelial protein expression and the tolerability of repeated doses of the PACE vehicles described here supports further investigation for protein supplementation applications in the lung.

For effective mucosal vaccination, mRNA targeting to APCs, which process and present antigen to other immune cells, is critical for vaccine efficacy (49). Due to the on-going global SARS-CoV-2 pandemic and high APC transfection with our platform, we chose to test the therapeutic potential of inhaled PACE-mRNA polyplexes as a mucosal vaccine against SARS-CoV-2. The rapid development of mRNA-based vaccines for SARS-CoV-2 was a historic achievement. To date, over 200 million US citizens have been vaccinated with Comirnaty or Spikevax. Not only were they the first vaccines to demonstrate safety and efficacy in reducing viral transmission, hospitalization, and death, but also the first mRNA-based therapeutics to gain FDA approval. Both vaccines target the S protein on the surface of the SARS-CoV-2 virus and deliver their mRNA cargo encapsulated in LNPs. These vaccines have the highest efficacy rates of all the vaccine platforms employed to combat the SARS-CoV-2 pandemic globally (50). Despite the remarkable safety and initial efficacy, declining immunity over time is a major limitation of these and other current vaccine platforms (51). Additionally, the continued emergence of new viral variants of concern (VOC) has further reduced vaccine efficacy (52-55). These limitations have spurred interest in new vaccination technologies that can combat the ongoing challenges posed by the SARS-CoV-2 pandemic.

Current mRNA vaccination strategies are focused on eliciting systemic immunity, primarily through the induction of IgG antibodies in serum and circulating antigen specific T cells (56, 57). However, growing evidence supports the superior effectiveness of vaccines that are delivered directly to the respiratory tract to combat respiratory viruses (58). The respiratory tract is the site of invasion and primary site of replication and disease manifestation for SARS-CoV-2 and its variants. Several studies investigating viral vector and protein-based vaccines have demonstrated that superior mucosal immunity can be achieved through direct antigen presentation across the mucosal surface of the lung (59-62). Mucosal vaccination strategies have been shown to improve upon systemic vaccination through the induction of tissue resident memory T and B cells, which can rapidly respond to viral invasion into the respiratory tract (31, 63). Additionally, mucosal vaccination can induce higher levels of antibodies in the respiratory mucosa (59). The induction of robust mucosal cellular and humoral immunity can more rapidly neutralize viruses upon entry into the respiratory tract, thereby preventing both infection and transmission (60).

Our PACE-mRNA vaccine induced both circulating and lung resident S specific CD8^+^ T cells, S specific memory B cells, and anti-S1 IgG in both serum and airways. The efficacy of PACE-mRNA as a mucosal vaccination strategy supports further investigation for respiratory viruses. Despite the impressive performance of approved mRNA-based vaccines and the potential to enhance protective immunity through mucosal delivery, as of the writing of this manuscript no mRNA-based vaccines have been reported to prevent SARS-CoV-2 infection following respiratory mucosal administration. This work is the first inhaled non-viral vector mRNA vaccine to demonstrate tolerability for repeated dosing and *de novo* induction of protective immunity against SARS-CoV-2.

## Conclusions

In summary, PACE-mRNA polyplexes can be formulated with blends of an end group-modified PACE and PACE-PEG to form small, consistent, and stable polyplexes. PACE polyplexes demonstrate favorable biocompatibility and protect mRNA from degradative enzymes. PEG content can improve polyplex characteristics (smaller size, more neutral surface potential, high mRNA loading) however dense PEG shells can interfere with transfection efficiency. Therefore, PEG content was optimized for lung specific delivery. Inhaled PACE-mRNA administration achieved high protein expression and effective lung targeting. Transfection occurred primarily in lung epithelial cells and APCs, two cell types which are highly relevant targets for a host of pulmonary diseases. Mucosal vaccination with PACE-mRNA induced systemic and lung resident adaptive immunity and protected mice from a lethal viral challenge.

## Acknowledgements

This work was supported by NIH grants UG3 HL147352 and R01 AI157488, the Fast Grant from Emergent Ventures at the Mercatus Center, and HHMI funding dedicated to collaborative research projects on SARS-CoV-2 and the disease it causes, COVID-19. A.S. is supported by the NIGMS T32GM136651. A.I. is an Investigator of the Howard Hughes Medical Institute. B.I. is supported by NIAID T32AI007517 and K08AI163493. T.M. is supported by NIAID T32AI007019. A.S.P. is supported by a K99/R00 Pathway to Independence award from the NIH (K99 HL151806) and a Postdoc-to-Faculty Transition Award from the Cystic Fibrosis Foundation (CFF; PIOTRO21F5).

## Author contributions

Conceptualization: AS, AI, WMS

Methodology: AS, MKG, HS, TM, BI, MR, JG, TL

Investigation: AS, MKG, HS, TM, BI, MR, JG, LA, TL, KS, ASP, RH

Visualization: AS, MKG, TM, BI

Funding acquisition: AI, WMS

Project administration: AI, HS, WMS

Supervision: AI, HS, WMS

Writing – original draft: AS, MKG

Writing – review & editing: AS, MKG, HS, TM, BI, MR, JG, LA, TL, KS, ASP, RH, AI, WMS

## Corresponding Authors

Correspondence and requests for data or materials should be addressed to H.S. or W.M.S

## Supplementary Materials

Materials and Methods Figs. S1 to S12

Table S1 to S2

