## Supplemental Materials for "Inhalable polymer nanoparticles for versatile mRNA delivery and mucosal vaccination"

Science
AAAS

### Supplementary Materials for

#### Inhalable polymer nanoparticles for versatile mRNA delivery and mucosal vaccination

Alexandra Suberi[1], Molly K. Grun[2], Tianyang Mao[3], Benjamin Israelow[3,4], Melanie Reschke[5], Julian Grundler[6], Laiba Akhtar[2], Teresa Lee[1], Kwangsoo Shin[1], Alexandra S. Piotrowski-Daspit[1], Robert J. Homer[7], Akiko Iwasaki[3,8], Hee Won Suh[1*], W. Mark Saltzman[1,2,9*]

**This PDF file includes:**

Materials and Methods
Figs. S1 to S12
Table S1 to S2

#### Materials and Methods

##### Materials and Reagents

Fetal bovine serum (FBS) was purchased from R&D Systems (Minneapolis, MN, USA). DMEM/F12 was purchased from Cytiva Life Sciences (Marlborough, MA, USA). Cleancap EGFP mRNA, Cleancap FLuc mRNA, and Cleancap Cre mRNA were synthesized by Trilink Biotechnologies (San Diego, CA, USA). Spike protein mRNA was extracted from Comirnaty vaccine obtained from pooled residual vaccine provided by Yale Health pharmacy. The Quant-iT RiboGreen RNA Assay Kit, Lipofectamine MessengerMAX, TrypLE Express, Phalloidin-Alexa Fluor 555, the Pierce BCA Protein Assay Kit, and Live/Dead Fixable Yellow Dead Stain Kit were obtained from Thermo Fisher Scientific (Walham, MA, USA).

*In vivo*-jetRNA was purchased from Polyplus-transfection (Illkirch, France). RediJect D-Luciferin Substrate was obtained from PerkinElmer (Waltham, MA, USA). Precellys hard tissue lysis tubes were obtained from Bertin Instruments (Montigny-le-Bretonneux, France). Glo Lysis Buffer and Bright-Glo luciferase substrate was purchased from Promega (Madison, WI, USA). Dispase was purchased from Corning (Corning, NY,USA). Collagenase II was purchased from Worthington Biochemical (Lakewood, NJ, USA). DNAse I was manufactured by Roche (Basel, Switzerland) and purchased through MilliporeSigma (St. Louis, MO, USA). 1,3-diamino-2-propanol was obtained from Alfa Aesar. Anhydrous dimethyl sulfoxide (DMSO), dichloromethane, and mPEG-OH (5k) were purchased from Sigma Aldrich. 1,1'-carbonyldiimidazole (CDI) was purchased from Acros Organics.

##### Polymer Synthesis and Characterization

PACE polymer (Mn = 4.0 kDa, Mw = 7.0 kDa, PDI 1.7) was synthesized according to previously reported procedures via terpolymerization of SA, MDES, and PDL or with additional mPEG-OH (5 kDa, Sigma Aldrich) for the synthesis of PACE-PEG (15). PACE end groups were activated by reaction with CDI following a modified literature procedure (16). Briefly, 40 eq. of CDI were dissolved in 10 mL dry DCM, and 1 g of PACE in DCM was added with stirring under argon. The resulting solution was stirred overnight at room temperature. The DCM layer was washed with 3 eq. DI water three times, and the DCM layer was collected, dried over $MgSO_4$, and dried under vacuum to obtain PACE-CDI, which was used as is without further purification.

For end group conjugation, 200 mg of PACE-CDI and 30 eq. of end group were dissolved in separate containers of 5 mL dry DMSO. Under argon atmosphere, the PACE solution was added dropwise into the amine solution with stirring in a cold bath (~8-10°C). The solution was stirred for 2 days at room temperature. At the end of reaction, 1.5 eq. of DCM was added to the reaction. Unreacted amine was extracted with 10 eq. of DI water, and the organic layer was washed three times with water. DCM was evaporated and the crude product was lyophilized overnight. Polymer was solubilized in THF and filtered through a 0.2 µm PTFE syringe filter and aliquoted into Eppendorf tubes. Solvent was evaporated under air and dried over vacuum overnight. Polymer was stored at -80°C until further use.

NMR samples were prepared in $d_1$-chloroform at a concentration of 10 mg/mL and recorded on an Agilent 400 MHz NMR spectrometer to check for completion of imidazole reaction with amine end groups by monitoring the peaks between 7-8.2 ppm, and to confirm the absence of free amine in the final product as well as the end group conjugation efficiency. Size exclusion chromatography was performed to confirm the molecular weight of the final polymer for transfection consistency. Measurements were performed on a ThermoFisher Ultimate 3000 UHPLC system with Malvern C–series column (CLM1031) connected to tandem detectors, Optilab® T-rEX refractive index detector and Dawn Heleos II light scattering detector (Wyatt Technology) at 50°C with DMF (0.1% w/v LiBr) at 0.8 mL/min. Samples were analyzed against PS standards (Agilent Easical PS-2 PL2010-0601). GPC samples were prepared by dissolving each polymer in 1 mL of 0.1% LiBr DMF overnight. Samples were filtered through 0.2 µm PTFE syringe filter before injection.

**Polyplex Production and Characterization**

All polyplexes were formed at a 100:1 or 50:1 weight ratio of polymer to mRNA. PACE and PACE-PEG were dissolved at 100 mg/mL overnight in DMSO (37°C, shaking). Prior to polyplex fabrication, PACE and PACE-PEG solutions were blended to produce a polymer mixture with the desired PACE-PEG concentration. mRNA and polymer were diluted into equal volumes of sodium acetate buffer (25 mM, pH 5.8) at the desired concentration. The polymer dilution was then vortexed for 15 s, mixed with the mRNA dilution, and vortexed for an additional 25 s. Polyplexes were incubated at room temperature for 10 min before use. To determine size, PDI, and surface charge, polyplexes were diluted to a final concentration of 50 µg/mL in deionized water and analyzed by dynamic light scattering (DLS, Zetasizer Pro, Malvern Analytical). mRNA loading was quantified using the Quant-iT RiboGreen RNA assay kit. The percentage of polymer associated mRNA was quantified by subtracting the unencapsulated mRNA (which produces a fluorescent signal) from a free mRNA control.

**PEG Surface Density and Conformation**

Freshly prepared polyplex NPs were diluted to six different concentrations (0.125, 0.25, 0.5, 0.75, 1, 2 mg/mL) and individually dialyzed against sodium acetate buffer using Slide-A-Lyzer dialysis cassettes (Thermo Scientific, 10 kDa MWCO) overnight. After dialysis, a sample aliquot of known volume was lyophilized to accurately determine the polyplex concentration. Subsequently, dialyzed samples and dialysate were directly injected (2 mL per injection) into a T-rEX refractive index detector (Wyatt Technology) at 0.3 mL/min *via* a syringe pump. The RI response of the dialysate was subtracted from the respective polyplex sample and the d*n*/d*c* value determined from the slope of the resulting linear fit.

The weight-averaged molar mass ($M_{w,SLS}$) was obtained by injecting nine concentrations (0.1, 0.2, 0.3, 0.4, 0.5, 0.6, 0.7, 0.8, 0.9 mg/mL) of 1 mL freshly prepared polyplex NPs into a Dawn Heleos II (Wyatt Technology) using a syringe pump with detector voltages normalized with dextran sulfate (15 – 20 kDa). The detector response was recorded using Astra 7 (Wyatt Technology) and global fitting based on Debye formalism was used to determine the NP molar mass.

The PEG surface density and conformation was calculated using a modified literature method (22) as follows, where $N_{PEG}$ represents the average number of PEG chains per polyplex NP, *SA* the NP surface area, $D$ the distance between PEG chains, $R_f$ the PEG Flory radius, and $DP$ the PEG degree of polymerization.

$$N_{PEG} = \frac{PACEPEG\ wt\% \times M_{w,SLS}}{M_{n,PACEPEG}}$$

$$SA = \frac{4}{3}\pi R_h^3$$

$$PEG\ density\ (\sigma) = \frac{N_{PEG}}{SA}$$

$$D = 2(\pi\sigma)^{-\frac{1}{2}}$$

$$R_f = (\frac{M_{n,PEG}}{DP})^{\frac{3}{5}} * 0.35\ nm$$

**In Vitro mRNA Uptake, Transfection, and Protection**

A549 cells were grown in a 1:1 mixture of DMEM/F12 supplemented with 10% fetal bovine serum (FBS) and 50 µg/mL gentamicin in a 37°C incubator under 5% $CO_2$. Twenty-four hours before mRNA delivery, cells were seeded at 50,000 cells/well in a 24-well plate and grown until 80-90% confluent. Immediately before delivery, the media was refreshed, and cells were treated with 1 µg/mL EGFP mRNA using PACE-E14 polyplexes or Lipofectamine MessengerMAX as a control. After 24 hours, cells were prepared for analysis by flow cytometry or fluorescence microscopy. For flow cytometry analysis, cells were washed once with sterile PBS, dissociated with TrypLE Express, and stained for live dead discrimination with the Zombie NIR Fixable Viability Kit following the manufacturer's instructions. Cells were then washed twice and analyzed by flow cytometry (Attune NxT). For microscopy, cells were washed three times with sterile PBS and fixed in 4% paraformaldehyde for 10 min. Cells were then permeabilized with 1% Triton X in PBS (15 min) and stained with 125 µg/mL phalloidin-Alexa Flour 555 in PBS with 1% bovine serum albumin (BSA, 60 min). Finally, cells were stained with 1 µg/mL Hoechst in PBS (10 min) and analyzed by fluorescence microscopy (Olympus LCPlanFl 20X, Olympus IX71). For RNase degradation experiments, transfection of EGFP mRNA-loaded PACE polyplexes with or without coincubation of RNase A was measured in HEK293T cells by flow cytometry. Polyplexes were incubated with RNase A (0.1 mg/mL; Thermo Fisher) for 10 minutes at 37 ºC, followed by incubation with Proteinase K (2 mg/mL; Thermo Fisher) for 15 minutes at 55 ºC. To validate the activity of the RNase enzyme, a formaldehyde denaturing gel was loaded with either pure mRNA or mRNA incubated with RNase A and Proteinase K as described above and the intensity of the mRNA band was visualized with SYBR Gold using a ChemiDoc MP imaging system.

**In Vitro Cytotoxicity Studies**

*In vitro* cytotoxicity of PACE-E14 polyplexes was assessed with the neutral red assay, following a previously published method (64). A549 cells were plated at 10,000 cells/well in a 96-well plate and grown overnight. The next day, the cell media was refreshed, and cells were treated with mRNA-loaded PACE-E14 polyplexes (0.125, 0.25, 0.5, 1, 2, 4 µg/mL mRNA). As a positive control, cells were treated with Lipofectamine MessengerMAX at the mRNA-to-reagent ratio recommended by the manufacturer. After 24 hours, cells were washed twice with PBS and treated with 100 µL neutral red working solution (40 µg/mL neutral red in complete medium) for two hours. Cells were then washed once with PBS, 150 µL of de-stain solution (50% ethanol, 49% water, 1% acetic acid) was added to each well, and the plate was vigorously mixed with an orbital shaker for 10 min. Finally, 100 µL of the solution was transferred to a fresh plate, and the absorbance was measured at 540 nm.

**Mice**

All procedures were performed in accordance with the guidelines and policies of the Yale Animal Resource Center (YARC) and approved by the Institutional Animal Care and Use Committee (IACUC) of Yale University and Yale Environmental Health and Safety. For reporter models of *in vivo* mRNA delivery, animal procedures were performed in a BSL-2 facility. Male BALB/c and Ai14 mice, age 7-20 weeks, were purchased from The Jackson Laboratory (Bar Harbor, ME, USA). BALB/c mice were used for luciferase expression assays, Cy5 labelled mRNA distribution studies and histology. Ai14 mice were used for flow cytometry experiments and lung microscopy. For vaccination experiments, B6.Cg-Tg(K18-ACE2)2Prlmn/J (K18-hACE2) mice were purchased from The Jackson Laboratory and bred and housed at Yale University. Female mice aged 8-12 weeks were used for all vaccinations.

**In Vivo mRNA Delivery in Reporter models**

For *in vivo* experiments, PACE-mRNA polyplexes were formulated at a final concentration of 0.1 mg/mL mRNA. For IT instillation, mice were anesthetized under 3% isoflurane (Patterson Veterinary), suspended by their incisors, the tongue was retracted, and 50 µL polyplex solution was pipetted into the posterior oral cavity. The tongue was held in the retracted position for the duration of 10 breaths. For IN delivery, mice were anesthetized under 3% isoflurane, the mouth was held closed and 50 ul polyplex solution was pipetted onto the nares of the animal. The mouth was held closed until no visible liquid droplet remained on the nares, approximately 5 breaths. For the control group, 5 µg of mRNA was delivered with *in vivo*-jetRNA following the manufacturer's instructions. For naked mRNA delivery, mRNA was diluted in sodium acetate buffer to a final concentration of 0.1 mg/ml mRNA and delivered by the usual IT method. For *in vivo* imaging (IVIS Spectrum, Perkin Elmer), mice were injected intraperitoneally with RediJect D-Luciferin Substrate 10-15 min before images were taken. To quantify luciferase expression, mice were euthanized, the heart was perfused with 15 mL PBS, and lung tissue was removed. The lungs were minced, transferred to 2 mL Precellys hard tissue lysis tubes with 1 mL Glo Lysis Buffer, and homogenized at 6500 rpm twice for 30 s (Precellys 24). The samples were then centrifuged at 20,000 x g for 10 min to pellet debris, and the supernatant was transferred to a fresh tube. 20 µL of tissue lysate was combined with 100 µL Bright-Glo luciferase substrate and luminescence was measured with an integration time of 10 s (Promega GloMax 20/20). Total protein concentration was quantified using the Pierce BCA Protein Assay Kit following the manufacturer's instructions.

For analysis by microscopy, mice were euthanized and the lungs were inflated with a 1:1 mixture of PBS and OCT before removal. A 3-mL syringe with a blunt-tip 20-gauge needle was inserted into the trachea, and a short length of suture was tied gently around the trachea. 1-1.5 mL of PBS/OCT was slowly injected into the lungs until fully inflated, the syringe was removed, and the lungs were quickly tied off. The tissue was then flash-frozen in OCT and sliced into 10 µm sections. Sections were stained with DAPI to visualize nuclei and analyzed by fluorescence microscopy.

For preparation of single cell suspensions for flow cytometry, lungs were inflated with 50 U/mL dispase and tied off with suture following euthanasia and perfusion. The tissue was minced and transferred to 5 mL of digestion buffer: 5 mg/mL Collagenase II and 1 mg/mL DNase I in PBS. The tissue was incubated shaking at 37°C for 30 min. After incubation, the digestion buffer was drawn up several times into a 5 mL syringe through an 18-gauge needle to break up remaining tissue pieces. On the final aspiration, the sample was discharged onto a 70 µm filter placed over a 50 mL conical tube. The syringe plunger was used to gently disrupt remaining tissue samples, then the filter was washed with 2 mL PBS with 0.5% BSA. The resulting single-cell suspension was pelleted (5 min, 1000 x g) and resuspended in 2 mL ACK lysis buffer to remove red blood cells. After 4 min, the buffer was neutralized with 8 mL of 10% FBS in PBS, strained once more through a 70 µm filter, pelleted (5 min, 1000 x g), and resuspended in FACS buffer (2% BSA in PBS). Cells were stained with anti-CD16/32 (TruStain FcX, S17011E, AB_2783137, BioLegend #156603) and for live/dead discrimination using the Live/Dead Fixable Yellow Dead Stain Kit (ThermoFisher L34959) following the manufacturer's instructions. Cells were stained with anti-CD45 (FITC, 30-F11, AB_312972 BioLegend #103107), anti-CD31 (Pacific Blue, 390, AB_10613457, BioLegend #102421), anti-CD326 (APC, G8.8, AB_1134105, BioLegend #118213), anti-CD11c (BV421, N418, AB_10897814, BioLegend #117329), and anti-CD11b (PerCP-Cy5.5, M1/70, AB_893233, BioLegend #101227) in FACS buffer for 30 min on ice, washed 2 times, and fixed with 4% PFA. Data was aquired on an LSR II BD instrument and analyzed with FlowJo software.

**In Vivo Histology**

PACE-E14 polyplexes with 10% PACE-PEG were formulated to a final concentration of 10 mg/mL polymer and 0.1 mg/mL mRNA, and 50 µL was delivered intratracheally to the lung. After 48 h, mice were euthanized, the heart was perfused with 5 mL of PBS, and the lungs and trachea were exposed. A blunt-tip needle was inserted into the trachea, and 1.5 mL of low molecular weight agarose (0.5% in PBS) was slowly injected to inflate the lungs. The trachea was tied off with suture, and the lungs were fixed for 48 h in 4% paraformaldehyde. Tissue samples were then embedded in paraffin, sliced to a thickness of 5 µm, and stained with hematoxylin and eosin (H&E). Representative sections were analyzed by a pulmonary pathologist blinded to the experimental groups.

**SARS-CoV-2 infection**

Animal procedures for SARS-CoV-2 infected mice were performed in a BSL-3 facility. Virus was obtained by infecting Vero E6 cells expressing hACE2 and TMPRSS2 with SARS-CoV-2 isolate hCOV-19/USA-WA1/2020 (NR-52281) as previously described (59, 65, 66). Mice were anesthetized under 3% isoflurane and $6x10^5$ PFU SARS-CoV-2 in 50 ul was pipetted directly onto the nares.

**mRNA Extraction from Comirnaty**

mRNA was extracted from the vaccine formulation with a TRIzol/chloroform separation method, as previously described (59). Briefly, aliquots of vaccine were dissolved in TRIzol LS (Thermo Fisher Scientific) at 1:6.6 vaccine to TRIzol. Following a 15 min incubation (37°C, shaking) 0.2

mL of chloroform was added per 1 mL of TRIzol. The solution was shaken vigorously for 1 min and then incubated at room temperature for 3 min. The solution was centrifuged at 12,000 x g for 8 min at 4°C. The aqueous layer containing the isolated mRNA was further purified with a RNeasy Maxi Kit purchased from Qiagen (Germantown, MD, USA) following the manufacturers protocol. The RNA was eluted from the column on the final step with sodium acetate buffer (25 mM, pH 5.8) warmed to 37°C. RNA purity and concentration was determined by Nanodrop measurement.

**Immunophenotyping vaccinated K18-hACE2 mice**

For vaccination experiments, K18-hACE2 mice between 6-8 weeks were immunized with a 10 µg dose of S loaded PACE-mRNA by IN delivery for a prime (day 0) and boost (day 28). Two weeks after boost delivery (day 42) immunophenotyping experiments were performed. On day 42, mice were intravenously injected with APC/Fire 750 CD45 Ab (30-F11, AB_2572116, BioLegend, #103154) 3 min before being euthanized. For CD8 T cell analysis, cells were stained with either anti-CD103 (BV421, 2E7, AB_2562901, BioLegend #121422), anti-CD3 (BV605, 17A2, AB_2562039, BioLegend #100237), anti-CD44 (BV711, IM7, AB_2564214, BioLegend #103057), anti-CD8a (PerCP/Cy5.5, 16-10A1, AB_2566491, BioLegend #305232), anti-CD69 (PE/Cy7, H1.2F3, AB_493564, BioLegend #104512), and PE SARS-CoV-2 S 539-546 MHC class I tetramer (H-2K(b)) for 30 min at 4°C. For B cell analysis, cells were stained with ant-GL7 (Pacific Blue, GL7, AB_2563292, BioLegend #144614), anti-IgM (BV605, RMM-1, AB_2563358, BioLegend #406523), anti-CD138 (BV711, 281-2, AB_2562571, BioLegend #142519), anti-CD19 (BV785, 6D5, AB_11218994, BioLegend #115543), anti-IgA (FITC, polyclonal, AB_2794370, SouthernBiotech #1040-02), anti-B220 (PerCP/Cy5.5, RA3- 6B2, AB_893354, BioLegend #103236), PE-SARS-CoV-2 RBD tetramer, anti-CD38 (PE/Cy7, 90, AB_2275531, BioLegend #102718), APC-SARS-CoV-2 RBD tetramer, and anti-IgD (AF700, 11-26c.2a, AB_2563341, BioLegend #405730) for 30 min at 4°C. For follicular helper T cell analysis, cells were stained with anti-CXCR5 (Biotin, L138D7, AB_2562126, BioLegend #145510) 69 min at 4°C, followed by staining with anti-PD1 (PE/Cy7, 29F.1A12, AB_10689635, BioLegend #135216), anti-CD44 (FITC, IM7, AB_312957, BioLegend #103006), anti-CD4 (AF700, GK 1.5, AB_493699, BioLegend #100430), anti-CD3 (BV421, 17A2, AB_2562553, BioLegend #100228), BV605 Streptavidin (BD Horizon, #563260) for 30 min at 4°C. Cells were washed with PBS and fixed with 4% PFA. Data was acquired on an Attune NxT Flow Cytometer and analyzed with FlowJo Software and gating was performed as previously described (59) (Fig. S12).

**Spike protein specific antibody measurements**

ELISAs were performed as previously described (29, 59, 65). Briefly, 96-well MaxiSorp plates (Thermo Scientific #442404) were coated with 50 µL/well of recombinant SARS-CoV-2 S1 protein (ACRO Biosystems S1NC52H3) at a concentration of 2 µg/mL in PBS and incubated overnight at 4°C. Plates were blocked with PBS containing 0.1% Tween-20 and 5% milk powder for an hour at RT. Serum or BALF was diluted in PBS with 0.1% Tween-20 and 2% milk powder and 100 µL per well was added for two hours at RT. Plates were washed five times and 50 µL of HRP anti-mouse IgG (Cell Signaling Technology #7076, 1:3,000) or HRP anti-mouse IgA (Southern Biotech #1040-05, 1:1,000) was added to each well for 1 hour. Plates were washed three times with PBS-T, 50 µL of TMB Substrate Reagent Set (BD Biosciences #555214) was added for 15 min and the reaction was stopped by the addition of 50 µL 2 N sulfuric acid. Plates were read at a wavelength of 450 nm and 570nm, and the difference plotted.

**Statistical Analysis**
Results were analyzed using GraphPad Prism (version 9.2.0 for Windows). Data are presented as mean ± SD. One- and two-way ANOVA with Dunnett's multiple comparison test or Tukey's multiple comparison were used where appropriate, unless otherwise indicated. Values were considered significantly different at $p < 0.05$.

**Graphical illustrations**
Created with Biorender.com

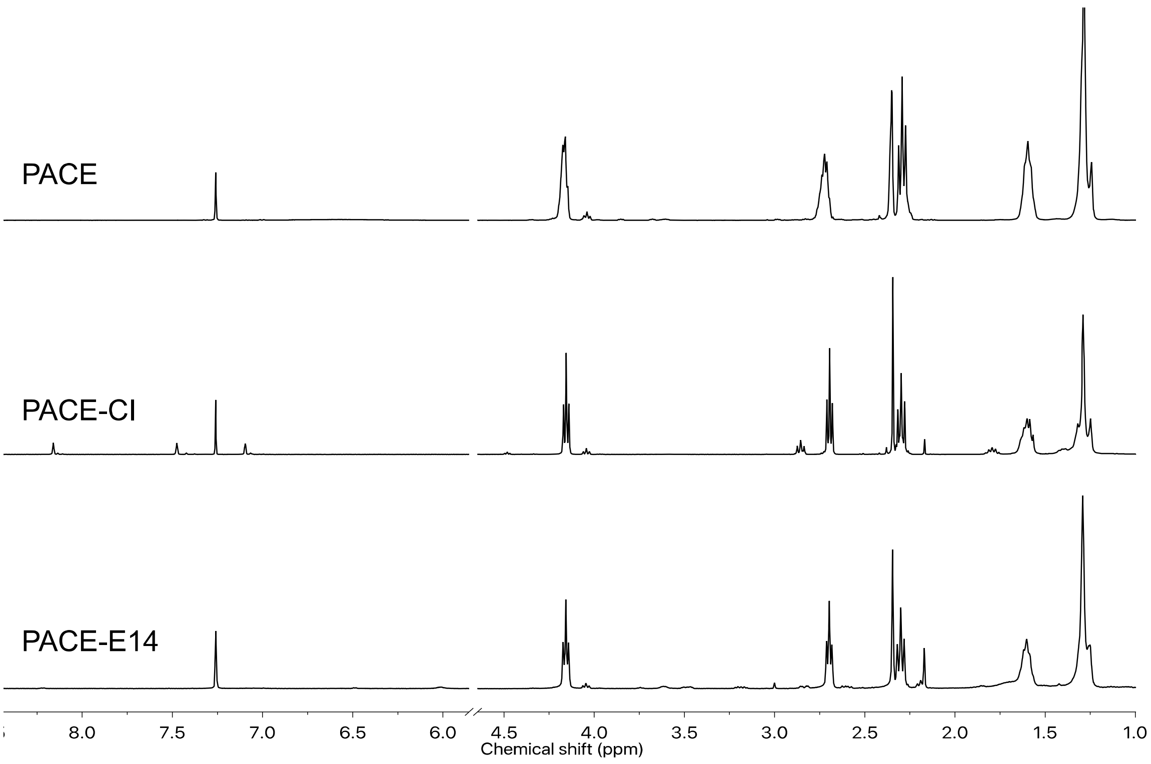


**Fig. S1.**
$^1$H NMR spectra of PACE, CDI reacted PACE (PACE-CI), and subsequent E14 reacted PACE (PACE-E14).

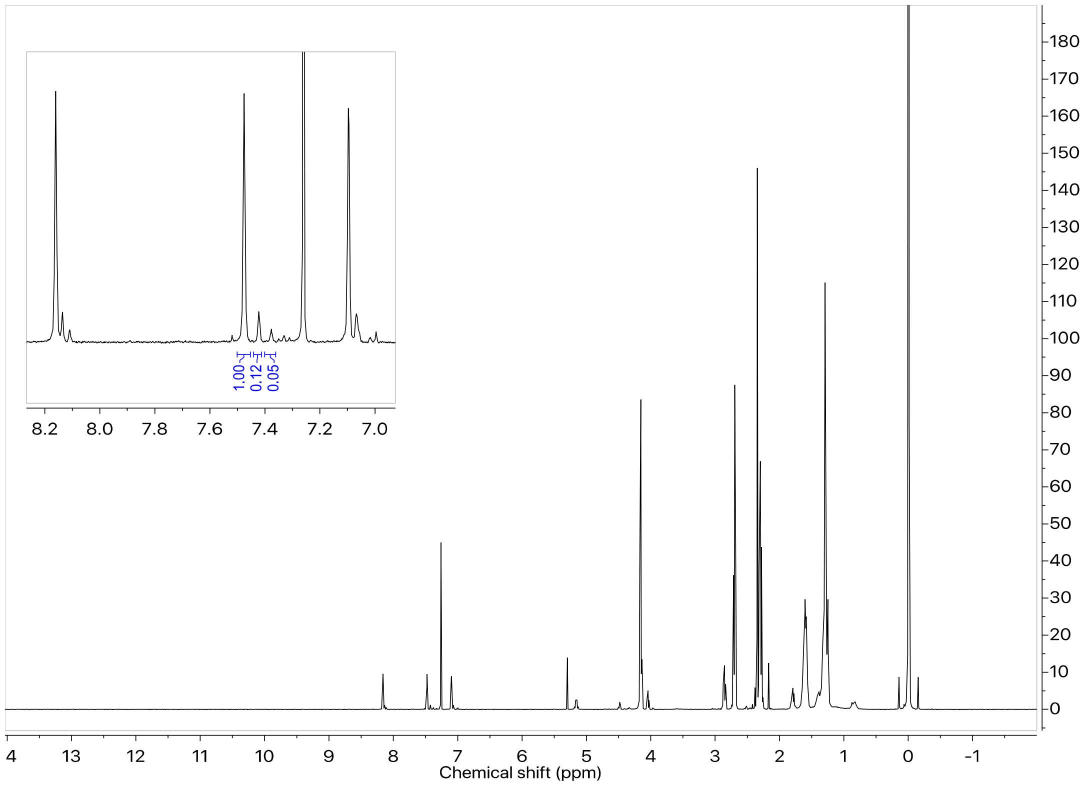


**Fig. S2.**
$^{1}$H NMR of PACE-CI. Imidazole peaks are shown in the top left corner with integrations representing the ratio of SBA to MDEA to PDL end-groups of the PACE base polymer, confirmed by $^{1}$H-$^{13}$C HSQC and HMBC (not shown). This indicates that PACE is primarily acid (SBA) ended (85%).

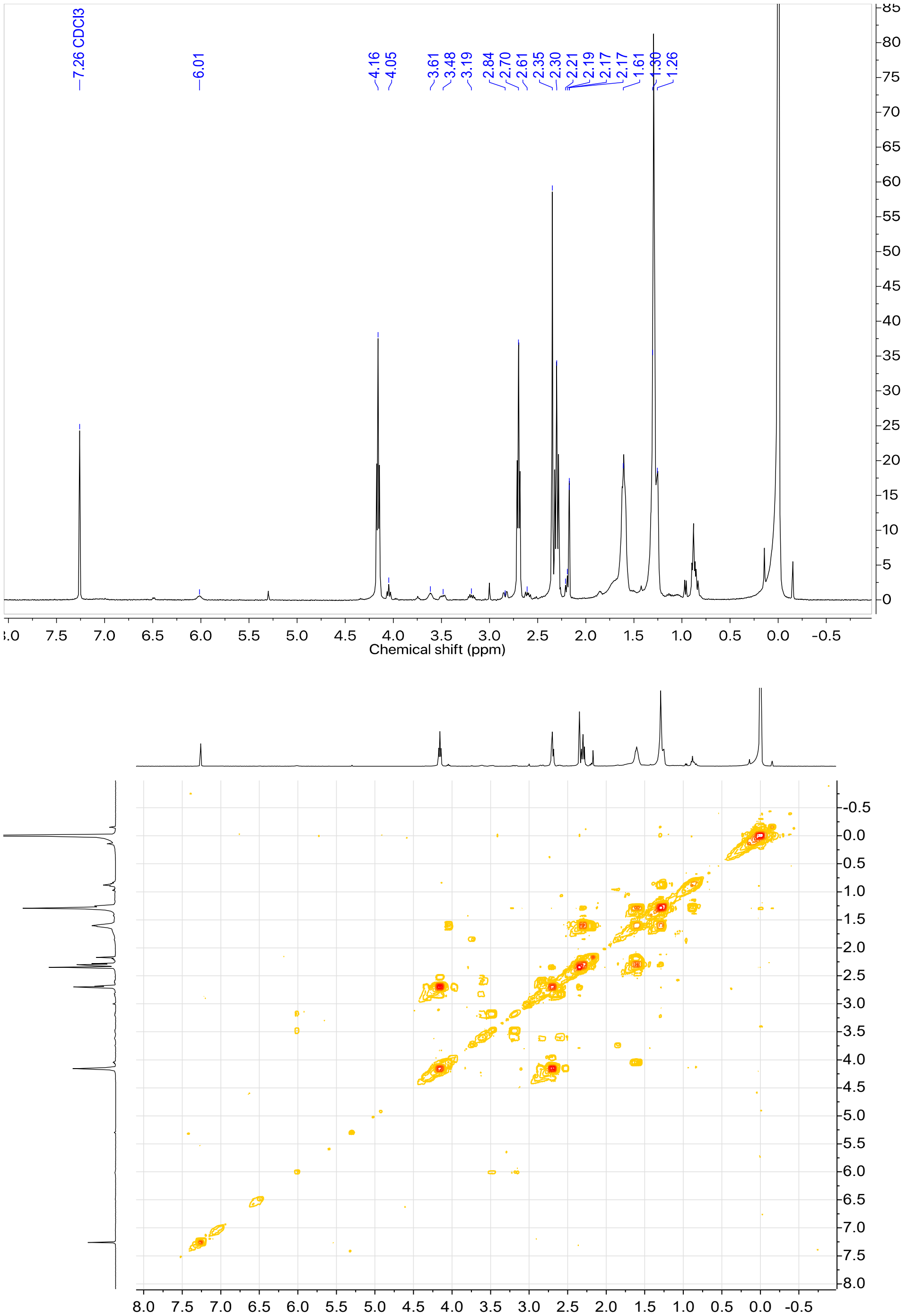

**Fig. S3.**
$^1$H NMR (top) and $^1$H-$^1$H COSY (bottom) spectra of PACE-E14. E14 end-group peaks are shown between 2.5-3.8 ppm.

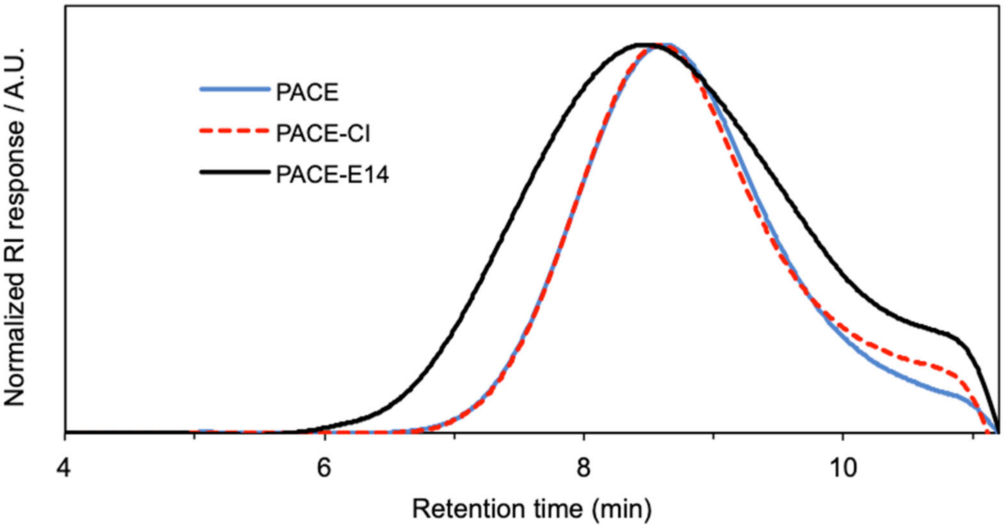


**Fig. S4.**
Overlay of GPC chromatograms of PACE, PACE-CI, and PACE-E14.

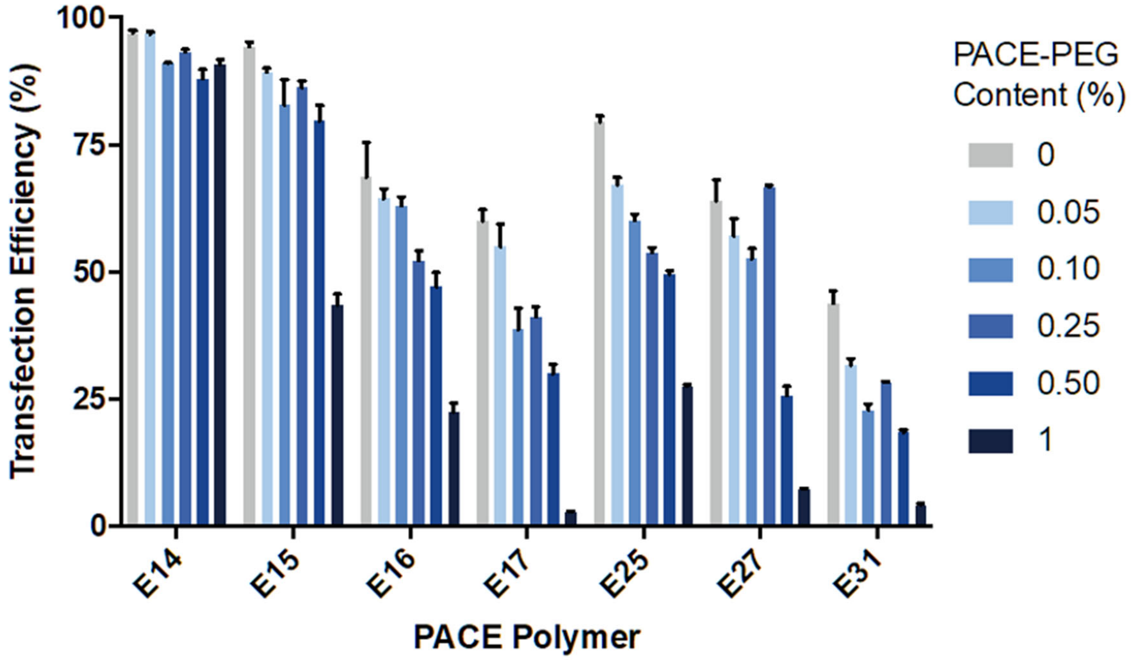


**Fig. S5.**
*In vitro* mRNA transfection with various EGFP mRNA PACE-endgroup polyplexes with varying PACE-PEG content in A549 lung epithelial cells by flow cytometry.

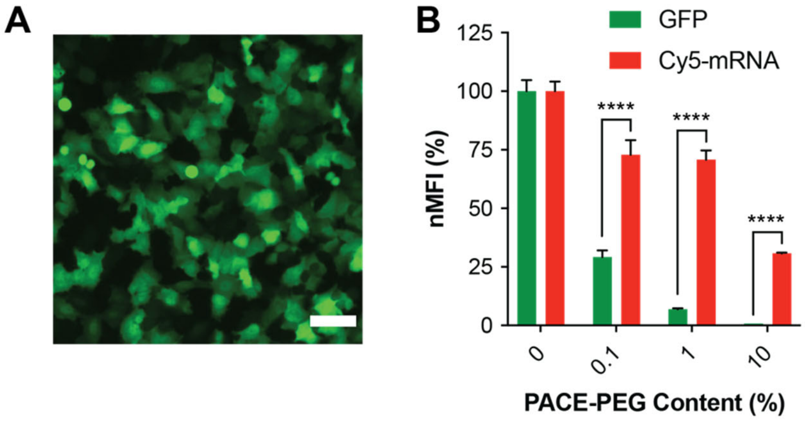


**Fig. S6.**
*In vitro* mRNA uptake and transfection with PACE-E14 polyplexes in A549 lung epithelial cells. **(A)** EGFP expression of EGFP mRNA delivered with non-PEGylated PACE-E14 polyplexes (scale bar, 75 µm). **(B)** Comparison of mRNA uptake and transfection efficiency using PACE-E14 polyplexes with varying PACE-PEG content. Mean fluorescence was normalized to non-PEGylated levels and compared by two-way ANOVA with Šidák's multiple comparison test. **** $p \leq 0.0001$.

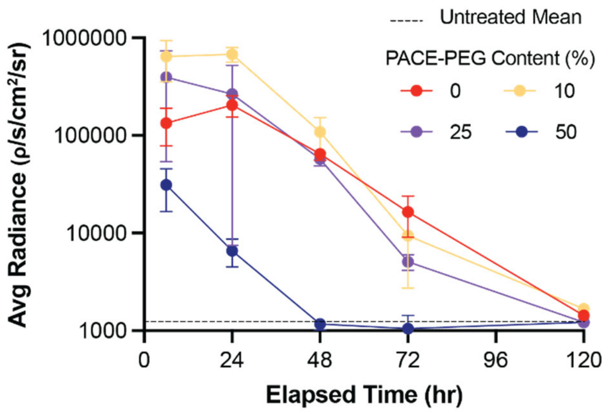


**Fig. S7.**
Kinetics of luciferase protein expression after IT delivery of 5 ug FLuc mRNA PACE-E14 polyplexes measured by IVIS.

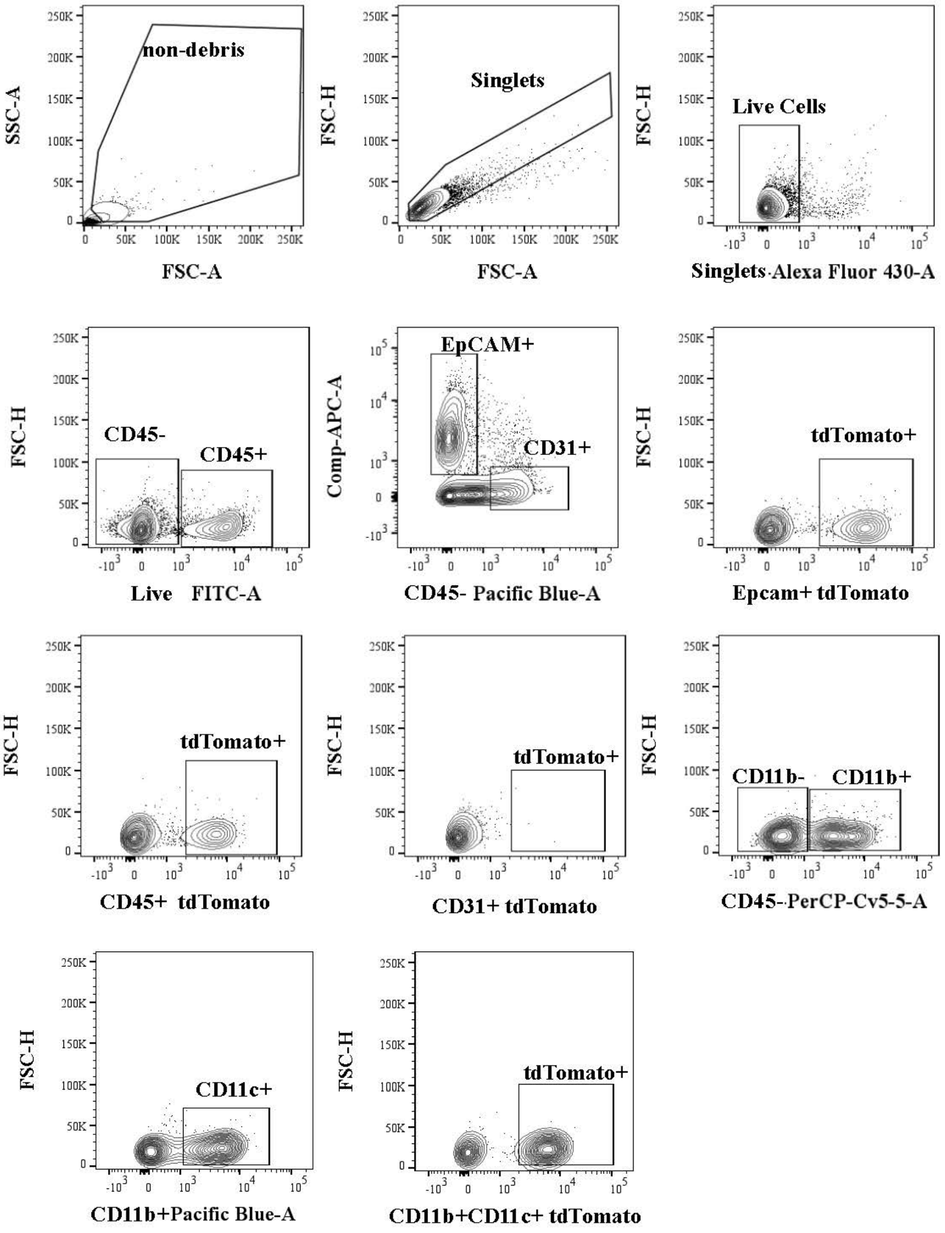


**Fig. S8.**
Gating strategy for Ai14 cells

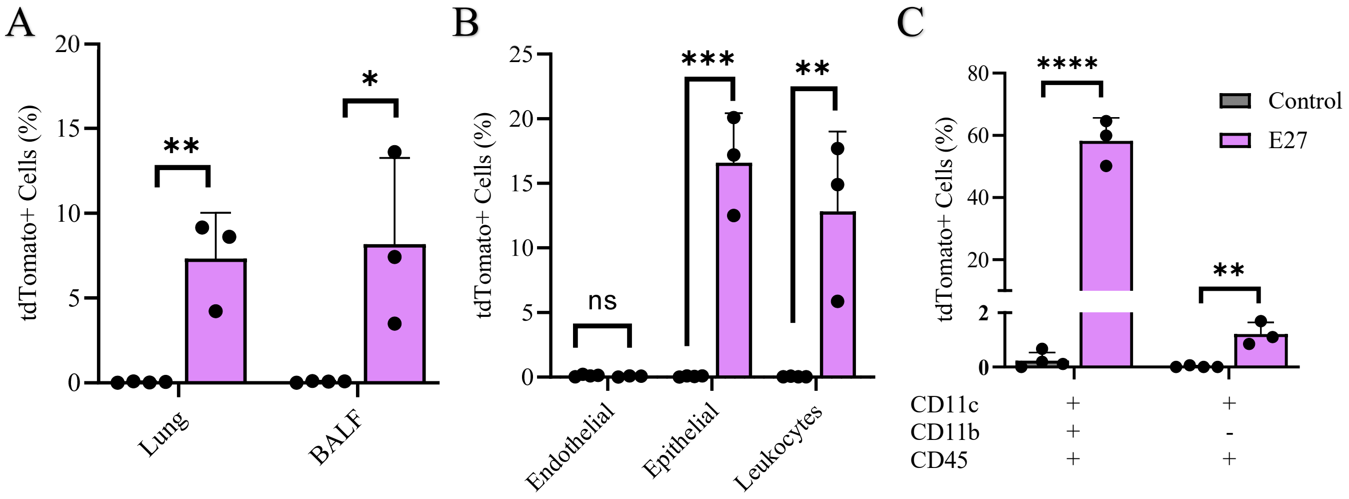


**Fig. S9.**
Percent of (**A**) all live cells in the lung and BALF, (**B**) endothelial, epithelial, or leukocyte cells in the lung and (**C**) APCs in the lung that express tdTomato 24 hours after delivery of PACE-E27 polyplexes (10% PACE-PEG) loaded with Cre mRNA. Statistical significance was calculated by multiple unpaired t-test with holm-sidak method. * $p \leq 0.05$, ** $p \leq 0.01$, *** $p \leq 0.001$, **** $p \leq 0.0001$.

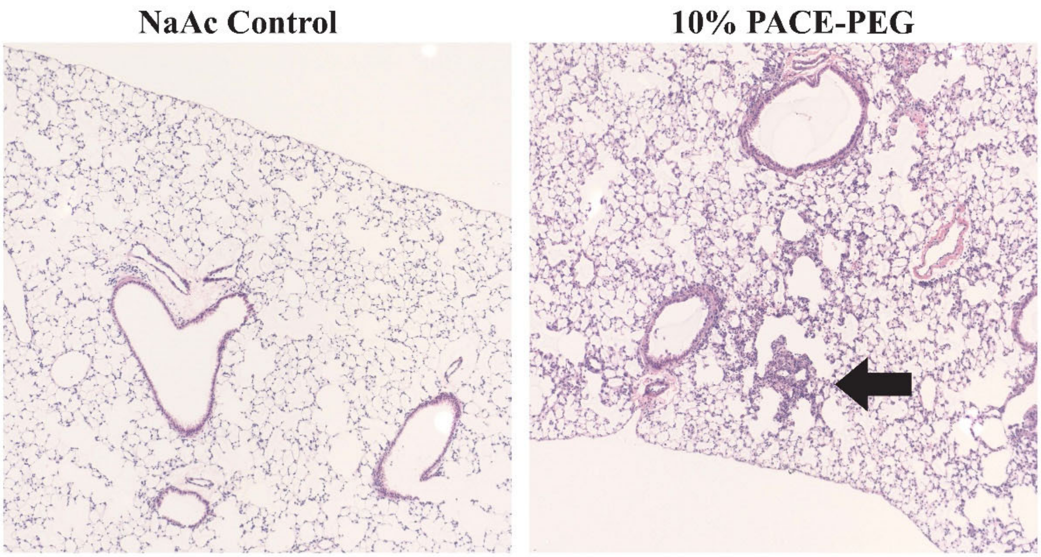


**Fig. S10.**
Lung histology 48 hours after inhaled mRNA delivery with PACE-E14 polyplexes demonstrates no evidence of necrosis or acute airway epithelial change. A representative image showing an area of focal pneumonitis following PACE-E14 polyplex delivery is shown (black arrow).

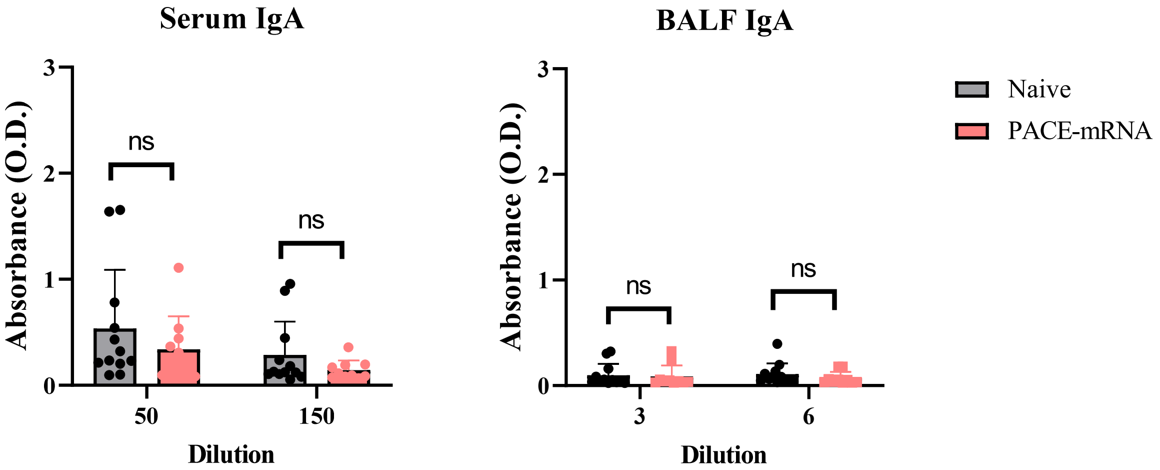


**Fig. S11.**
Serum and BALF analysis of anti-S1 IgA levels measured by ELISA in naïve (untreated) and PACE-mRNA vaccinated mice.

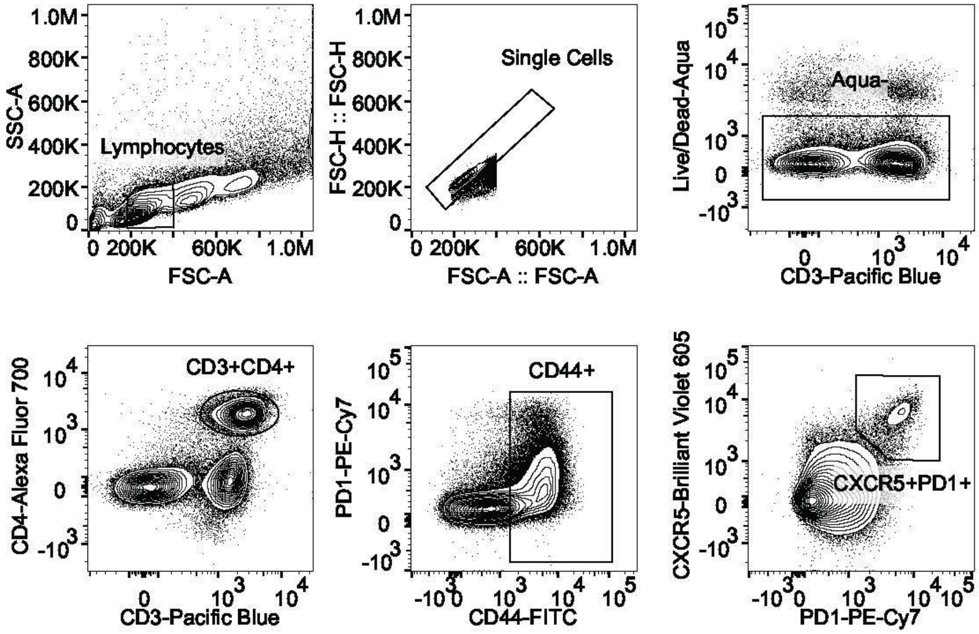


**Fig. S12.**
Gating strategy for $T_{FH}$ cells.

| Polymer | Mn (kDa) | Mw (kDa) | PDI |
|---|---|---|---|
| PACE | 4.0 | 7.0 | 1.74 |
| PACE-CI | 4.1 | 7.7 | 1.86 |
| PACE-E14 | 4.1 | 9.5 | 2.30 |
| PACE-PEG | 12.2 | 13.9 | 1.14 |

**Table S1.** Summary of polymer characterization by GPC.

| PACE-PEG (%) | PEG density (PEG/100 $nm^2$) | SD | $D/R_f$ | SD | $R_g/R_h$ | Rf/D |
|---|---|---|---|---|---|---|
| 1 | 0.1 | 0.06 | 8.86 | 3.13 | 1.03 | 0.13 |
| 10 | 0.3 | 0.09 | 3.52 | 0.50 | 1.06 | 0.29 |
| 25 | 1.2 | 0.33 | 1.72 | 0.23 | 1.16 | 0.59 |

**Table S2.**

Results from polyplex PEG density calculations.
